# Alpha phase coding supports feature binding during working memory maintenance

**DOI:** 10.1101/2024.01.21.576561

**Authors:** Mattia F. Pagnotta, Aniol Santo-Angles, Ainsley Temudo, Joao Barbosa, Albert Compte, Mark D’Esposito, Kartik K. Sreenivasan

**Affiliations:** Helen Wills Neuroscience Institute, University of California Berkeley, Berkeley, CA, United States of America; Division of Science and Mathematics, New York University Abu Dhabi, Abu Dhabi, United Arab Emirates; University of Utah, Salt Lake City, UT, United States of America; Institut d’Investigacions Biomèdiques August Pi i Sunyer (IDIBAPS), Barcelona, Spain; Laboratoire de Neurosciences Cognitives et Computationnelles, INSERM U960, École Normale Supérieure - PSL Research University, 75005 Paris, France; Department of Psychology, University of California Berkeley, Berkeley, CA, United States of America; Center for Brain and Health, New York University Abu Dhabi, Abu Dhabi, United Arab Emirates

## Abstract

The ability to successfully retain and manipulate information in working memory (WM) requires that objects’ individual features are bound into cohesive representations; yet, the mechanisms supporting feature binding remain unclear. Binding (or swap) errors, where memorized features are erroneously associated with the wrong object, can provide a window into the intrinsic limits in capacity of WM that represent a key bottleneck in our cognitive ability. We tested the hypothesis that binding in WM is accomplished via neural phase synchrony and that swap errors result from perturbations in this synchrony. Using magnetoencephalography data collected from human subjects in a task designed to induce swap errors, we showed that swaps are characterized by reduced phase-locked oscillatory activity during memory retention, as predicted by an attractor model of spiking neural networks. Further, we found that this reduction arises from increased phase coding variability in the alpha-band over a distributed network of sensorimotor areas. Our findings demonstrate that feature binding in WM is accomplished through phase coding dynamics that emerge from the competition between different memories.

**Significance:** We investigate the neural basis of working memory, focusing on how feature binding is accomplished and how binding or ‘swap’ errors arise. Using magnetoencephalography, we found that stable phase-locking of alpha oscillations supports correct feature binding, while swap errors correlate with reduced alpha phase preservation, localized to specific brain areas. These findings align with a biologically-plausible computational model predicting that temporal synchrony in neuronal firing underpins feature binding. This work advances our understanding of the neural mechanisms of working memory, providing empirical support for theories of time-based binding and demonstrating the utility of biophysically-realistic models in human neuroimaging studies.

## 1. Introduction

Working memory (WM) is the ability to retain and manipulate information when it is no longer present in our environment, which introduces the possibility for abstract concepts and plans to influence our behavior ^1–4^. The well-described capacity limits of WM ^5–7^ hence represent a key constraint on higher order cognition such as cognitive control ^8^. In real life, WM capacity is not just dependent on the maintenance of individual visual features (e.g., color, orientation, spatial location), but also of the conjunctions or bindings between them ^9–12^. Thus, understanding how information is bound in WM is of critical importance to understand the intrinsic limits of this executive function.

One proposed mechanism for feature binding in WM is temporal synchrony between populations of neurons that encode individual features ^13,14^. This idea relies on the notion of temporal or phase coding—that is, the two different neuronal populations encoding the information of distinct features of a conjunction fire at the same phase of an ongoing oscillation, enabling these features to be bound together in WM ^15–17^. While the idea of phase coding is broadly consistent with empirical evidence of synchronized oscillatory dynamics within or between different brain regions during WM ^18–23^, a biologically plausible computational implementation of phase coding has been elusive ^24,25^ and direct evidence for phase coding has been lacking. A recent model ^26^ provides a plausible neural architecture of the cortico-cortical circuits that implement feature binding in WM, in which phase coding emerges from the lateral inhibition between the different neuronal populations that store information about distinct (competing) conjunctions ^27^. In the present study, we sought to test a central prediction of this model: that disruptions in oscillatory phase should be associated with misbinding or ‘swap’ errors ^26^, where an inaccurate response to the target item is accurate relative to a non-target item (e.g., if a subject shown a red square and a blue circle mistakenly reports the color of the circle as red) ^10,11^. If this prediction were validated, it would provide a key empirical demonstration that phase coding serves as an organizational principle during WM maintenance.

We tested our hypothesis using magnetoencephalography (MEG) recordings collected from human subjects performing a task designed to induce swap errors. Our results showed a characteristic within-trial phase-locking in the alpha-band during WM maintenance. In parietal-occipital sensors contralateral to the visual stimuli, the consistency over trials of such alpha phase-locking was reduced in swap trials compared to high-performance on-target trials. Importantly, these effects did not generalize to other WM errors, suggesting that such deterioration in phase-locked oscillatory activity is a hallmark of swaps. To understand why phase relationships are compromised in swap trials, we considered a measure of variability in the instantaneous frequency of alpha oscillations and showed that swaps are characterized by increased variability. We further localized these effects in contralateral areas in premotor, motor, parietal, and visual cortices. These results suggest that during WM maintenance feature binding is accomplished via alpha phase coding, while swaps are produced by unstable phase-locked activity in distributed sensorimotor areas.

## 2. Results

### 2.1. Identification of swap errors

We analyzed MEG data previously collected from 26 human subjects who performed a delayed-response task designed to induce swaps ^28^ (Fig. 1). Subjects were instructed to remember a briefly shown lateralized display of 3 circles (stimulus presentation: 0.2 s). After a brief memory retention interval (delay period: 2 s), subjects reported the location of each of the circles, which were sequentially cued by their color (report period: self-paced) in a random order. Each subject completed a total of 500 trials of this WM task (see section 4.2).

**Fig. 1.**
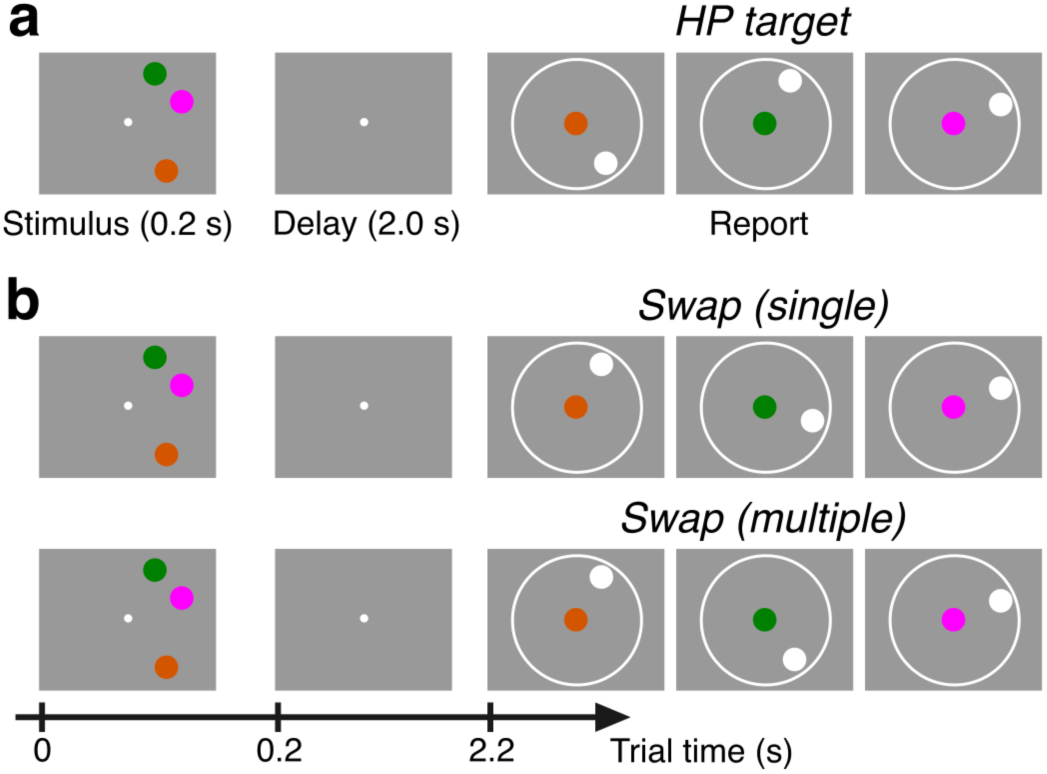
Experimental design. a–b. Schematic representation of the temporal structure of each trial, with examples of high-performance (HP) response (a) and swap errors (b). In the schematic depiction of the report period, the colored circles represent the cues, while the white circles represent the subject’s responses. In b, the examples depict trials in which the location of one (upper) or more (lower) of the non-cued circles are mistakenly reported.

We used continuous measures of response errors to model subjects’ behavior. First, we evaluated the histograms of response deviations from target feature value (i.e., cued item) and from all non-target feature values (i.e., uncued items; Fig. 2a). Since we enforced a minimum distance between the feature values in each trial (15° of polar angle, see section 4.2), the histogram of response deviations from non-target features shows a dip around zero (Fig. 2a). This may obscure the presence of a central peak, which would indicate the presence of swaps. To correct for these effects of minimum feature distance, we used an approach which consisted in subtracting the distribution expected in the absence of swaps from the observed non-target deviations ^29^. The distribution of non-target response deviations corrected for the effects of minimum feature distance revealed that the responses were clustered around the non-target features, with the histogram showing a central peak (Fig 2a), which indicates the presence of swaps in our experiment. We employed a maximum likelihood approach to distinguish high-performance (HP) trials (location of all circles reported accurately; Fig. 1a) from swap trials (location of one or more of the non-cued circles mistakenly reported; Fig. 1b), and from low-performance (LP) trials (location of one or more circles reported inaccurately) based on the subjects’ responses ^30,31^ (see section 4.2). After categorizing the trials into HP trials, LP trials, and swaps, we separately analyzed the histograms of response deviations by trial type. As expected, the histogram of response deviations from the target feature showed a central peak for each trial type, with the distribution being more concentrated for the trials identified as HP trials than for LP trials and swaps (Fig. 2b). On the other hand, the histograms of response deviations from non-target features, both observed and those corrected for the effects of minimum feature distance, showed a central peak only for the trials identified as swaps (Fig. 2c–d), confirming the correct identification of trial types.

**Fig. 2.**
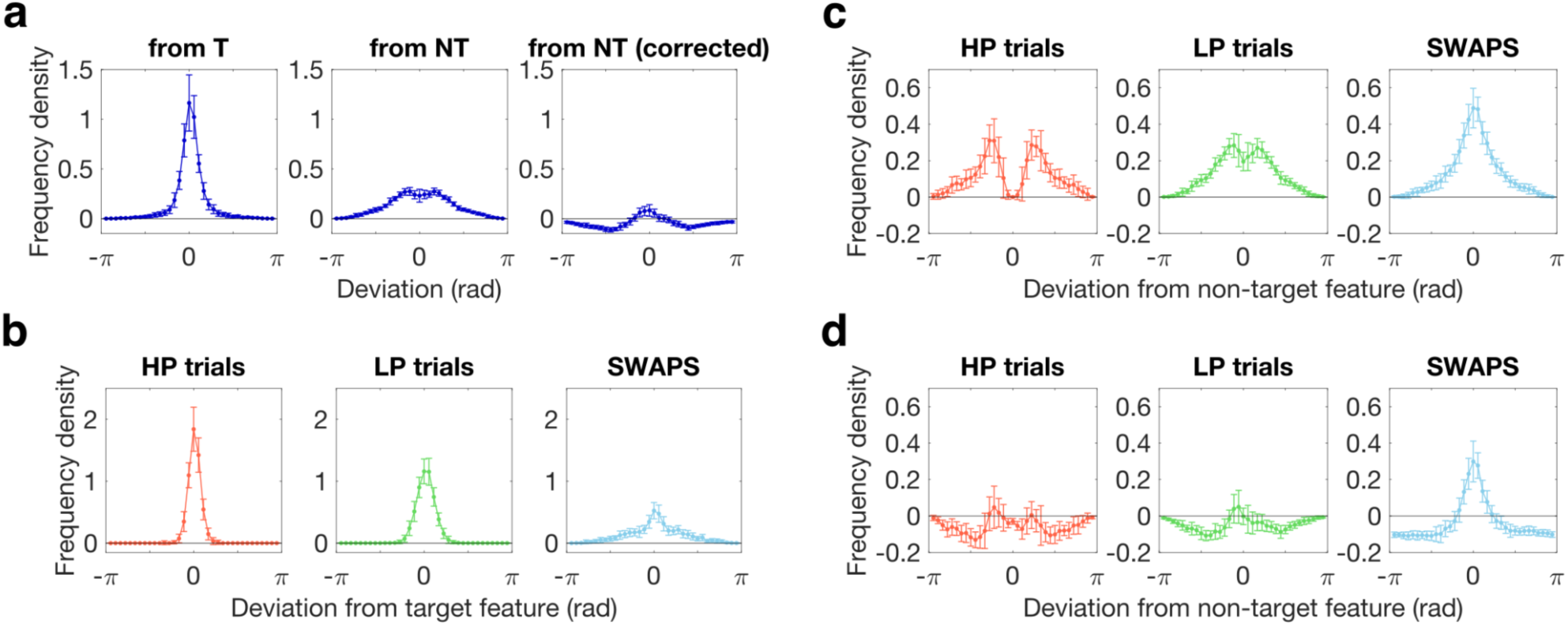
Distribution of response errors. **a** Histograms of response deviations across all subjects. The first plot (*left*) shows the distribution of response deviations from the target feature (T). The second plot (*middle*) shows the distribution of response deviations from the non-target features (NT). The third plot (*right*) shows the distribution of NT response deviations corrected for the effects of minimum feature distance. **b** The histograms of response deviations from the target feature (in radians) are shown for HP trials (red), LP trials (green), and swaps (blue). **c** The histograms of response deviations from the non-target features (in radians) are shown for HP trials (red), LP trials (green), and swaps (blue). **d** Same as in c, but showing the distributions after correction for the effects of minimum feature distance. In each figure, the error bars represent the mean and standard deviation across subjects, using bins with a width of 0.1745 rad (i.e., 10° of polar angle), in the range between [-π, π] rad (i.e., [-180°, +180°]).

We observed all three trial types in all subjects: HP trials ranged from 16–439 across subjects (*M*=92.81, *SD*=88.06); swaps ranged from 22–196 (*M*=116.38, *SD*=47.79), and LP trials ranged from 44–328 (*M*=198.73, *SD*=64.03). In a subsequent analysis (see section 2.4), we distinguished between trials that had one (Fig. 1b, upper example) or multiple (Fig. 1b, lower example) swap reports. Memory performance for each trial type was assessed using the von Mises concentration parameter, which is a measure of the concentration of the distribution of response errors and provides a proxy for WM precision (see section 4.2). A paired samples *t*-test was performed to evaluate whether there was a difference in memory precision, as measured by the concentration parameter, between HP trials (*M*=23.93, *SD*=8.12, range 14.47–52.56), swaps (*M*=11.37, *SD*=4.82, range 4.98–26.02), and LP trials (*M*=11.00, *SD*=3.58, range 5.15–21.32). Unsurprisingly, the results indicated that memory was more precise for HP trials as compared to swaps (*t*(25)=13.4808, *p*<0.0001) and LP trials (*t*(25)=11.1340, *p*<0.0001). Performance was comparable for swaps and LP trials (*t*(25)=0.7953, *p*=0.4340), indicating that these two trial types were equated for overall performance, but crucially differ in feature binding alone.

### 2.2. Less stable phase-locked dynamics in swaps during WM maintenance

Our hypothesis entailed that HP trials are characterized by stable phase-locked activity during WM maintenance, while swap trials are induced by noisy fluctuations in the phase of oscillatory activity related to the maintenance of individual features. This prediction was motivated by a recent computational model ^26^ that employs two ring-attractor networks—one for each feature space (here, color and spatial location)—to explicitly simulate the independent storage of item features. Briefly, the model implements feature binding through the selective synchronization of bump pairs across the two networks (for full model details, see Ref. 26). Color-location associations are maintained through correlated oscillatory activity, relying on phase coding ^32^. Swaps occur when there are abrupt changes in the phase relationship between oscillating bumps, while stable phase coding dynamics support correct feature binding. To measure changes in phase relationships in our MEG data, we used the Phase Preservation Index (PPI) ^33^, which captures for each time point the consistency over trials of the within-trial phase differences with respect to a reference time (in our case, the memory delay onset at t=0.2 s; Fig. 1)—that is, their level of phase clustering in polar space ^34^. We selected this reference time point following the approach used in previous work on the attractor model ^26^. Thus, PPI provides a measure of the consistency of frequency-specific local phase-locking over trials, as a function of time. We estimated PPI for each MEG sensor over time points in the delay period (0.2–2.2 s; Fig. 1) and frequencies 1–50 Hz separately for each trial type, and then compared PPI between HP trials and swaps. Since PPI is sensitive to the number of trials, we performed the analysis by equating the number of HP and swap trials (see section 4.5). To leverage the lateralized nature of neural signals, we used a pooling procedure that aligned the MEG sensors on the left side with the hemisphere contralateral to stimulus presentation, and the sensors on the right side with the ipsilateral hemisphere (see section 4.3). Behavioral performance (mean absolute error) and the proportion of trial types did not vary significantly between left-hemifield and right-hemifield trials (all *p*-values>0.19).

PPI was higher in HP targets than swaps in the alpha-band. We found a cluster in the observed data, extending over frequencies between 8–14 Hz and latencies around ∼0.5 s after delay onset (Fig. 3a). PPI differences between trial types were most pronounced over 25 parietal-occipital sensors, contralateral to stimulus presentation (Fig. 3b), with medium effect size (Cohen’s *d* in the range 0.170–0.576 across sensors in the observed cluster). The cluster-based permutation test revealed that PPI was significantly higher in HP targets compared to swaps (*p_perm_*=0.0160), with phase preservation reduced for swaps in the alpha-band, showing a steeper decrease than HP targets (Fig. 3c), as predicted by model simulations of synchronization-based feature binding ^26^. In order to test the specificity of the observed alpha-band effects, we calculated the z-scores of PPI over frequencies for each time point in the delay, separately for each trial type. We found positive z-scores at low frequencies in the delta/theta range (up to ∼6 Hz) and negative z-scores at higher frequencies in the beta/gamma range (above ∼15 Hz), while the middle frequencies in the alpha-band (∼10 Hz) were characterized by positive z-scores that were higher than those at adjacent frequencies, both low and high (Fig. 3d). These results reveal that while there is a general tendency for PPI to decrease faster at higher frequencies than lower frequencies, alpha deviates from this trend.

**Fig. 3.**
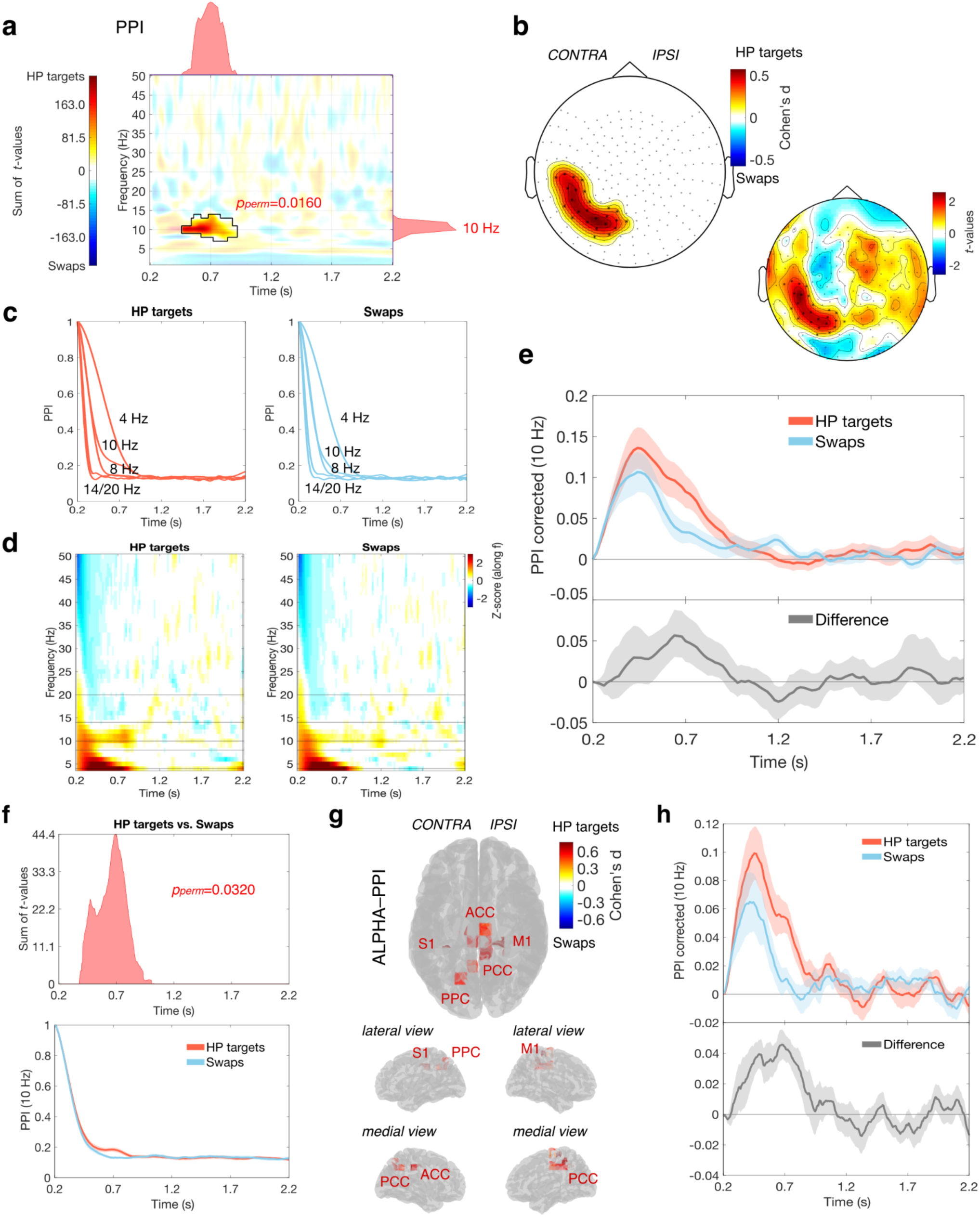
PPI differences between HP targets and swaps. **a** Time-frequency distribution of the sum of PPI differences across MEG sensors (*t*-values). The black contour highlights the statistically significant cluster. The marginal plots on the right and on top represent respectively the time-collapsed frequency distribution and frequency-collapsed time distribution of the differences between trial types. **b** Topography plot with superimposed effect sizes of PPI differences between HP targets and swaps, for each MEG sensor of the observed cluster (*t*-values are shown on the smaller topography plot on the right). **c** Time course of PPI in HP targets (left) and swaps (right) at different frequencies, highlighted by the black horizontal lines in the next panel. **d** Z-scores along frequencies for each time point in the delay, in HP targets (left) and swaps (right). **e** Time course of PPI corrected at 10 Hz for HP targets (red) and swaps (blue), together with the difference between the two (gray). The red and blue shadings (top) represent the standard error of the mean, while the gray shading (difference; bottom) represents 95% confidence intervals (CIs). CIs were estimated using the bias-corrected and accelerated method on a bootstrap distribution of across subjects differences, obtained by resampling with replacement 10,000 times. **f** Time distribution of the sum of 10 Hz PPI differences between HP targets and swaps, across source points in source-space (*t*-values; top), and time course of the average PPI across source points of the positive cluster for each condition (bottom). The shadings represent the standard error of the mean. **g** Glass brain with superimposed effect sizes of the observed differences between HP targets and swaps. **h** Time course of PPI corrected at 10 Hz for HP targets (red) and swaps (blue) across source points of the cluster in the observed data, together with the difference between the two (gray; shadings are as in panel e).

To confirm that the observed PPI effects were due to within-trial phase synchronization in the alpha-band, we performed the following control analysis. We repeated the PPI estimation (including the procedure to equate the number of trials between HP targets and swaps – see section 4.5), but this time after randomly shuffling the signals from each MEG sensor over time, which disrupted any phase relationships present in the data. We then computed ‘PPI corrected’ estimates as the difference between PPI estimates from the main analysis and those obtained from the control analysis. The results obtained from shuffled data (i) showed that the rate of PPI reduction over time increases monotonically with frequency, including for the alpha-band, and (ii) confirmed the presence of a plateau/asymptote for PPI at all frequencies (∼0.13 on average across subjects; Supplementary Information, Fig. S1a–b). The PPI corrected results showed, instead, that phase synchronization takes place specifically at alpha frequencies in our WM task (Supplementary Information, Fig. S1c–d), and that HP targets are characterized by more stable dynamics relative to swaps during the delay period (Fig. 3e).

In a separate analysis, we compared PPI estimates between HP targets and LP trials and did not find any statistically significant differences (the lowest *p*-value from the permutation test among the identified clusters was *p_perm_*=0.187). Critically, given that memory performance on swaps and LP trials was comparable, this finding rules out the possibility that differences in alpha-band PPI are merely a function of task difficulty or generalized performance impairment, and instead suggests that our results are specific to feature binding errors. When we directly tested for differences between LP targets and swaps, we again found no statistically significant PPI differences (the lowest *p*-value from the permutation test among the identified clusters was *p_perm_*=0.554). To assess the direction of these non-significant effects, we examined PPI and PPI corrected estimates at 10 Hz for each trial type, across sensors in the previously observed cluster (see Fig. 3b), as well as the time course of the difference between trial types (Supplementary Information, Fig. S2). The results showed a gradient from HP targets to swaps in both PPI and PPI corrected estimates, with LP targets in the middle—characterized by lower phase-locked alpha activity than HP targets during the delay period (Supplementary Information, Fig. S2b–e), but more stable than swaps (Supplementary Information, Fig. S2c–f).

To identify the underlying cortical sources exhibiting differences in local phase-locked activity between HP targets and swaps, we used a source reconstruction technique to localize MEG sources of activity, and repeated the PPI analysis in source-space (see section 4.7). PPI estimation was here restricted to the frequency of interest of 10 Hz, which was selected as the peak of the time-collapsed frequency distribution of observed PPI differences in sensor-space (see Fig. 3a). In source-space, we observed a positive cluster (higher PPI in HP targets than swaps) in the data (Fig. 3f), which included 17 source points localized in the cingulate cortex and cortical areas contralateral to stimulus presentation (Fig. 3g). Effect size peaked in source points localized in the ventral anterior cingulate cortex–ACC and dorsal posterior cingulate cortex–PCC, ipsilateral primary motor cortex–M1, and contralateral areas in primary somatosensory cortex–S1 and precuneus/posterior parietal cortex–PPC (see Table 1). The cluster-based permutation test identified that there was a significant difference in 10 Hz PPI between HP targets and swaps (*p_perm_*=0.0320). Like in the sensor-space analysis, the PPI corrected results showed that alpha phase-locked activity is characterized by more stable dynamics in HP targets than swaps during the delay period (Fig. 3h).

**Table 1.** PPI differences between HP targets and swaps: peak effect sizes.

| Anatomical region | MNI coordinates (x, y, z) | Cohen's *d* |
| --- | --- | --- |
| Ventral ACC | -17.5, -20, 42.5 | 0.694 |
| Dorsal PCC | 7.5, -44.5, 55 | 0.702 |
| M1 | 20, -32.5, 67 | 0.732 |
| S1 | -30, -32.5, 42.5 | 0.766 |
| Precuneus/PPC | -17.5, -69.5, 2.5 | 0.616 |

*Note.* ACC and PCC stand for anterior and posterior cingulate cortex, respectively. M1 stands for primary motor cortex. S1 stands for primary somatosensory cortex. PPC stands for posterior parietal cortex.

We note, however, that it is difficult to draw conclusions about the precise timing of effects due to the intrinsic limitations of the cluster-based permutation testing, which only provides an approximation of the effect extent ^35,36^. Further, PPI may not be best suited for determining the timings of the observed effects. Because it is equal to 1 at the reference time point and decays quickly over time ^26,33^, PPI estimates close to said reference are effectively inflated. This can be appreciated by examining the time course of PPI at different frequencies (e.g., in HP targets; Fig. 3c): estimates tend to decrease over time to an asymptotic value. Thus, PPI may be most sensitive to differences that are in the early delay period of the task.

To address these temporal biases in PPI estimation and provide a more complete picture of the timing of the observed PPI effects, we conducted a separate analysis in which we computed PPI using the middle of the delay as the reference time (*t_ref_*=1.2 s), rather than using the delay onset (see section 4.5). Similar to the results in Fig. 3, we observed a parietal-occipital cluster that displayed a trend toward alpha-band PPI increase in HP targets compared to swaps in posterior sensors, contralateral to stimulus presentation (Supplementary Information, Fig. S3a–b), though it did not reach statistical significance (*p_perm_*=0.0679). We again observed that PPI has a fast roll-off toward a plateau/asymptote (Supplementary Information, Fig. S3c) and the z-score analysis confirmed that alpha deviates from the general tendency for PPI, characterized by a faster decrease at higher than at lower frequencies (Supplementary Information, Fig. S3d). In addition, the results of the PPI corrected analysis confirmed that within-trial phase synchronization takes place specifically in the alpha-band (Supplementary Information, Fig. S3f–h). Overall, these findings confirm that phase synchronization is disrupted during maintenance in swap trials, but reinforce the notion that the timing of the PPI effects should be interpreted cautiously.

### 2.3. Increased phase coding variability in swaps

The attractor networks model ^26^ suggests that phase-locked dynamics allow maintaining color-location conjunctions in WM, but these phase-locked dynamics can be broken by abrupt noisy fluctuations, which lead to misbinding of memorized features and swaps. We confirmed the neurophysiological prediction of this model, in that trials containing swaps have a lower PPI compared to HP targets. However, PPI does not directly address the origin of instabilities in the phase of alpha oscillations (Fig. 4a). A degradation of alpha phase preservation could occur either as a result of fluctuations in alpha phase that continues throughout the delay period or as a result of transient fluctuations in alpha phase. To disambiguate these two scenarios, we calculated frequency sliding (FS), which captures the instantaneous temporal fluctuations in oscillation peak frequency (Fig. 4b)—that is, time-varying changes in the instantaneous frequency of the oscillator ^37^. In order to demonstrate the feasibility of using FS to capture the noisy fluctuations that induce swaps, we estimated FS from the signals obtained from the two attractor networks model ^26^. In Fig. 4c–d, we depict an example trial where a noisy fluctuation at around t=2.7 s reverses the internetwork correlation, ultimately producing a swap (Fig. 4e). Our results demonstrate that FS is able to capture the timing of these fluctuations (Fig. 4f). In the first network, FS varies suddenly at around t=2.7 s, accurately capturing the time when the internetwork correlation reverses. This effect is also visible on the difference in FS between the two networks (see black trace in Fig. 4f).

**Fig. 4.**
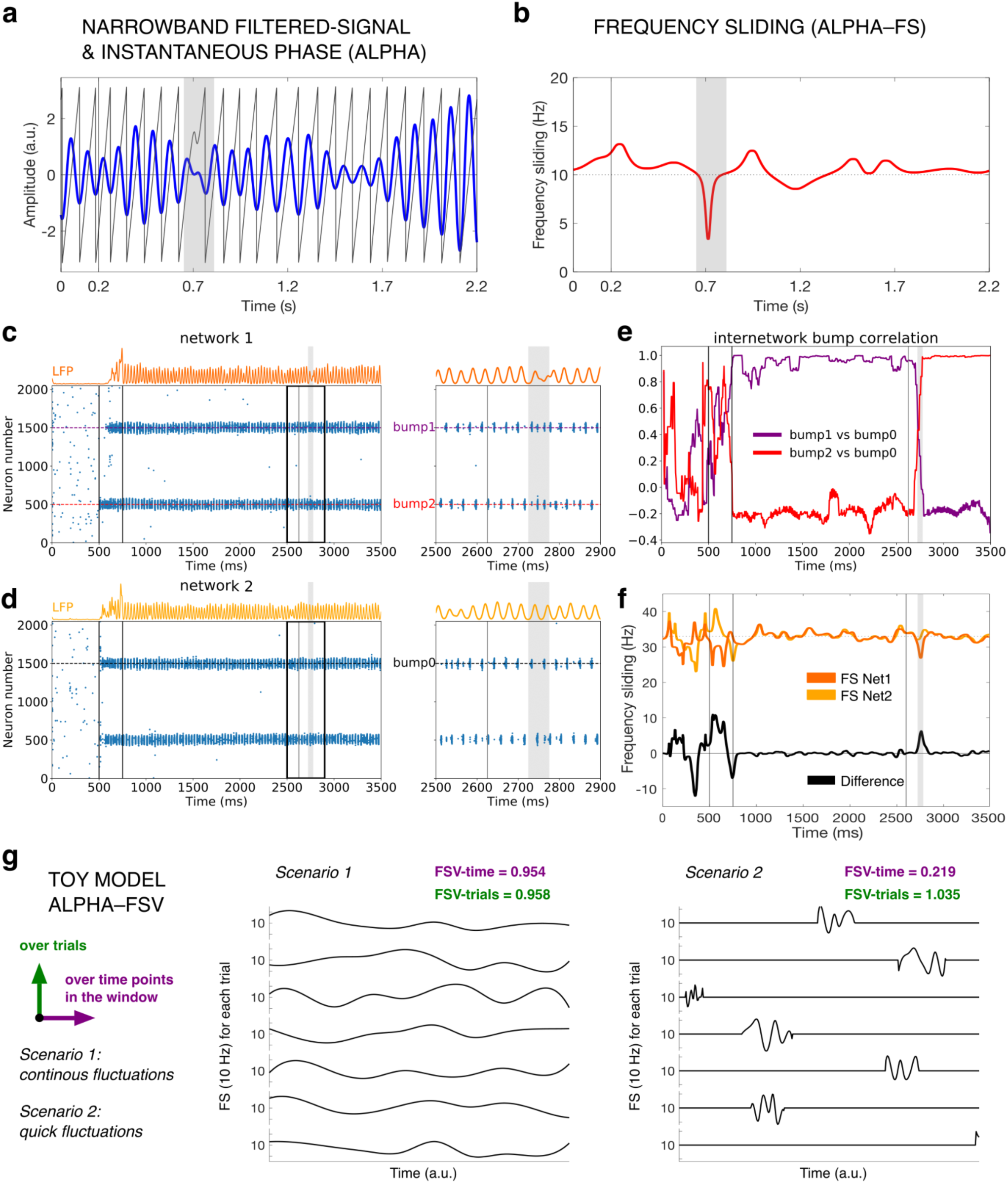
Instantaneous phase fluctuations, frequency sliding (FS), and frequency sliding variability (FSV). **a** Example from a swap trial of the narrowband filtered-signal in the alpha-band (in blue) and its instantaneous phase (in black) during the delay period. The signal was obtained from the first subject, from one of the contralateral parietal sensors where we observed significant PPI difference. **b** Single-trial FS estimated from the signal represented in panel a. In panels a–b, the gray shadings highlight a noisy fluctuation in the alpha phase and how this can be captured by FS. **c–d** Raster plots of a sample simulation ^26^ from the two ring attractor networks (left), together with the delay period zooms showing clear bump oscillatory activity (right). **e** In the simulated example, a noisy fluctuation reverses these correlations suddenly producing a swap trial, as shown by the internetwork bump correlation plot. **f** Frequency sliding (FS) estimated from the simulated field signals of the first (dark orange) and second network (light orange) at peak-frequency, from the model depicted in panels c–d. The difference in FS between the two networks is depicted in red (bottom). The vertical gray shading marks the time of a large, abrupt noisy fluctuation, leading to a swap error during the readout. **g** Toy model illustrating the estimation of alpha-FSV over time points or over trials, and alternative scenarios for how a reduction in alpha phase preservation in swaps is produced by noisy fluctuations in instantaneous alpha frequency with different temporal profiles.

Unlike PPI, FS is a reference-independent measure; thus, it overcomes the limitation of PPI discussed in section 2.2 to provide a better estimate of the timing of observed effects. This is a useful feature also because previous studies have recognized that, in addition to WM maintenance, encoding and recall are also noisy processes ^11^; consequently, swaps may result from a failure in correctly binding visual features during any of these three phases of WM ^38–41^. We selected a frequency of interest at the peak of the time-collapsed frequency distribution of PPI differences (10 Hz; see Fig. 3a), and derived FS at single-trial level. We then estimated a measure of ‘alpha-FS variability (FSV) over time’ in a sliding window during the time between the stimulus presentation and the delay period (0–2.2 s) by taking the median between trials of the standard deviation of FS over time points, separately for HP targets and swaps (see section 4.6). We also estimated the median standard deviation of FS over trials on the sliding window to derive a measure of ‘alpha-FSV over trials’, separately for HP targets and swaps (see section 4.6). The frequency sliding variability (FSV) provides a measure of variability in phase coding. In a perfect phase code, neurons can encode information through the timing of their spikes relative to a reference oscillation, such as a neural rhythm (e.g., alpha), and a neuron would always fire at the same phase for the same stimulus or state. However, in reality, there is some randomness or noise, and spike timing fluctuates. Phase coding variability quantifies how much this timing fluctuates. The implication is that higher phase coding variability might indicate the presence of noise or interference, making it harder to decode information. Lower phase coding variability implies a more reliable and efficient neural code, potentially allowing for better discrimination of stimuli or states, as predicted by the attractor model. Differences between HP targets and swaps in both measures of alpha-FSV would indicate that swaps are induced by noisy fluctuations that are more sustained over time (scenario 1 in Fig. 4g), while if the source of variability is predominantly over trials, this would suggest that noisy fluctuations in swaps happen quickly and more suddenly (scenario 2 in Fig. 4g and as predicted by the attractor model, Fig. 4f).

We did not find any statistically significant differences between HP targets and swaps in the alpha-FSV over time (the lowest *p*-value from the permutation test among the identified clusters was *p_perm_*=0.368). Alpha-FSV over trials, however, was significantly higher in swaps compared to HP targets (*p_perm_*=0.0180) in a cluster extending into the second half of the delay period (approximately between 1.182–2.045 s; Fig. 5a). The differences in alpha-FSV over trials were most prominent in 20 contralateral, parietal-occipital sensors (effect size ranging between *d*=0.088 and *d*=0.321 across cluster sensors; Fig. 5b). This and the previous PPI analysis represent convergent findings, as the groups of electrodes showing the biggest effects largely overlapped between the two analyses (compare Fig. 3b and Fig. 5b). These results appear to indicate that noisy fluctuations in swaps happen suddenly in time. To directly test whether this is the case, we compared the two measures of alpha-FSV over trials and alpha-FSV over time (see section 4.6). If the two measures are comparable, this would indicate that the variability in phase coding is induced by noisy fluctuations that are sustained over time (scenario 1 in Fig. 4g). On the other hand, if alpha-FSV over trials is significantly higher than alpha-FSV over time, phase coding variability is induced by more abrupt fluctuations (scenario 2 in Fig. 4g). A paired samples *t*-test was performed to evaluate whether there was a difference in swaps, between alpha-FSV over trials (*M*=1.29, *SD*=0.17, range 0.90–1.49) and alpha-FSV over time (*M*=0.84, *SD*=0.21, range 0.38–1.11). The results indicated that alpha-FSV over trials was significantly higher than alpha-FSV over time (*t*(25)=47.5922, *p*<0.0001; Supplementary Information, Fig. S4), suggesting that, in swaps, noisy fluctuations in the alpha oscillations’ peak frequency happen abruptly.

**Fig. 5.**
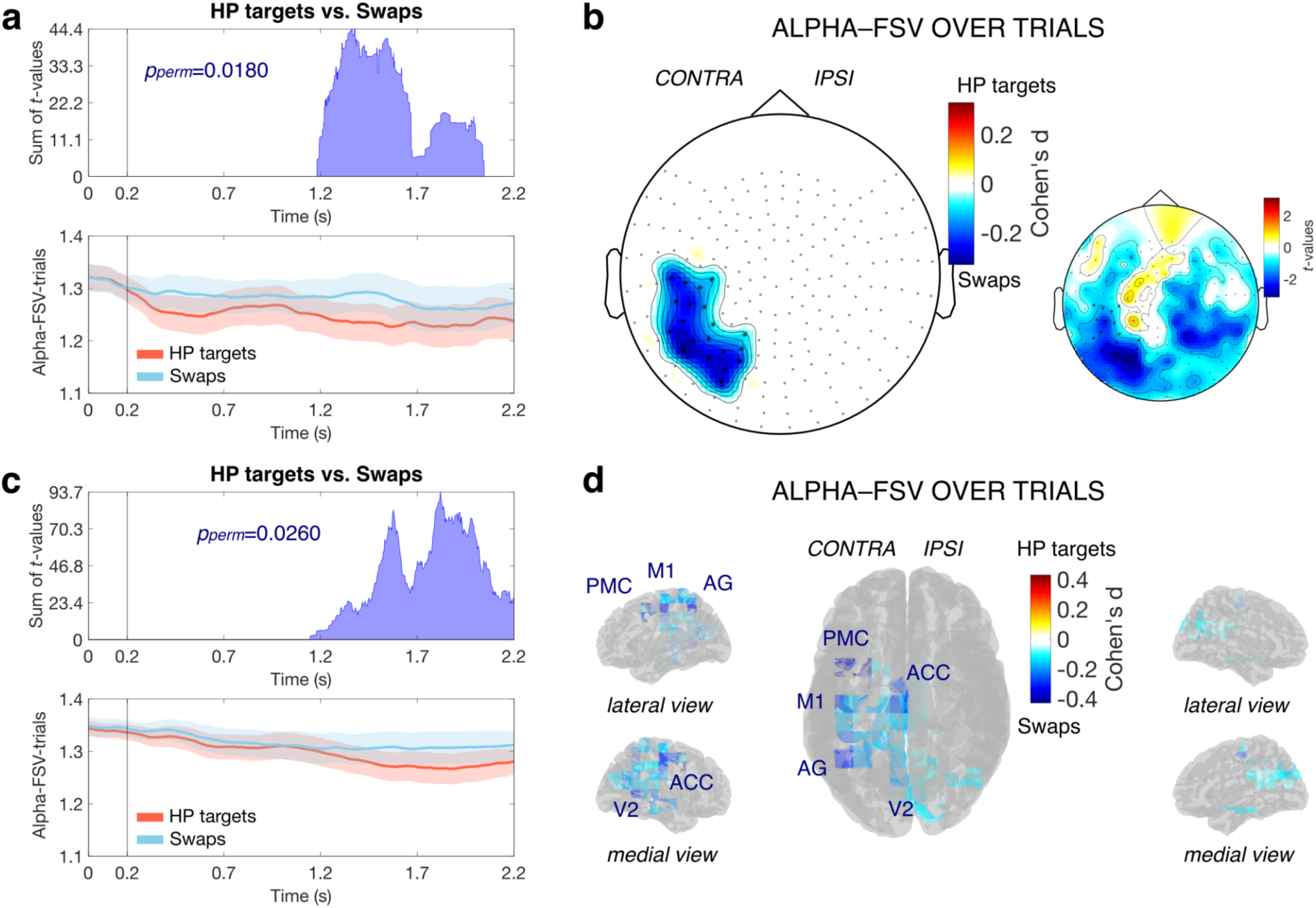
Differences in alpha-FSV over trials between HP targets and swaps. **a** Time distribution of the sum of alpha-FSV differences across MEG sensors (*t*-values; top), and time course of the average alpha-FSV across sensors of the cluster found in the data for each condition (bottom). The shadings represent the standard error of the mean. **b** Topography plot with superimposed effect sizes of the differences between HP targets and swaps, for each MEG sensor of the observed cluster (*t*-values are shown on the smaller topography plot on the right). **c** Time distribution of the sum of alpha-FSV differences across source points in source-space (*t*-values; top), and time course of the average alpha-FSV across source points of the observed cluster for each condition (bottom). The shadings represent the standard error of the mean. **d** Glass brain with superimposed effect sizes of the differences between HP targets and swaps.

As for the PPI, in a separate control analysis, we compared alpha-FSV over trials between HP targets and LP trials and did not find any statistically significant differences (the lowest *p*-value from the permutation test among the identified clusters was *p_perm_*=0.192). When we directly compared LP targets and swaps, we again found no statistically significant differences in alpha-FSV over trials (the lowest *p*-value from the permutation test among the identified clusters was *p_perm_*=0.815). To assess the direction of these non-significant effects, we examined alpha-FSV over trials for each trial type, across sensors in the previously observed cluster (see Fig. 5b), as well as the time course of the difference between trial types (Supplementary Information, Fig. S5). Our results showed a gradient from HP targets to swaps in alpha-FSV over trials, with LP targets in the middle—characterized by higher phase coding variability than HP targets during the delay period (Supplementary Information, Fig. S5b), but slightly lower than swaps (Supplementary Information, Fig. S5c).

Analogous to our PPI analysis, we repeated the FSV analysis in source-space (see section 4.7). The source-space results confirmed that alpha-FSV over trials was higher in swaps than HP targets. A negative cluster was found extending approximately over latencies 1.148–2.200 s (Fig. 5c) and 92 source points, localized for the most part in contralateral cortical areas (Fig. 5d). Effect size peaked in source points localized in premotor cortex–PMC, motor area M1, angular gyrus–AG, visual area V2, and anterior cingulate–ACC (see Table 2). The cluster-based permutation test indicated that there was a significant difference in alpha-FSV over trials between swaps and HP targets (*p_perm_*=0.0260). In line with the sensor-space results, we did not find any statistically significant differences in alpha-FSV over time (lowest *p_perm_*=0.833 among clusters). Together these results suggest that, in areas contralateral to stimulus presentation, correct feature binding in WM is supported by stable phase coding in the alpha-band, while swap errors are the result of its perturbations. In swaps, noisy fluctuations in the alpha oscillations peak frequency happen quickly and abruptly rather than in a more sustained fashion over time.

**Table 2.** Differences in alpha-FSV over trials between HP targets and swaps: peak effect sizes.

| Anatomical region | MNI coordinates (x, y, z) | Cohen’s *d* |
| --- | --- | --- |
| PMC | -30, 5, 55 | 0.430 |
| PMC | -42.5, 5, 42 | 0.386 |
| M1 | -42.5, -20, 55 | 0.361 |
| AG | -42.5, -57, 55 | 0.305 |
| V2 | -5, -57, 5 | 0.291 |
| ACC | -5, -20, 42 | 0.290 |

*Note.* PMC stands for premotor cortex. M1 stands for primary motor cortex. AG stands for angular gyrus. ACC stands for anterior cingulate cortex. V2 stands for secondary visual cortex.

### 2.4. Distinct possible sources of swaps

The term ‘swap’ is agnostic as to whether features are reciprocally/symmetrically exchanged between two memorized items (i.e., the two items are misbound to each other), or unidirectionally exchanged (i.e., only one item is misbound to a non-target) ^42^. That is, in a scenario where a subject is presented with a red item at location 1 and a green item at location 2, and, when asked to recall the location of the red item, erroneously reports location 2, it is unclear whether they would necessarily report location 1 when asked to recall the location of the green item. If so, this would imply a reciprocal/symmetric exchange of features. Here, we defined the trials with only one swap (i.e., a single non-target report) as ‘single-swap’ and the trials with multiple swaps as ‘reciprocal-swap’ (see Fig. 1b for examples). The nature of our task, which involved a full report of all memorized items, allows us to ask (i) whether swaps are necessarily reciprocal/symmetric, and (ii) if both reciprocal-swap and single-swap trials exist, whether phase coding variability differs between these two types of swaps, potentially implying a different underlying mechanism. Our analysis focused on FSV for two reasons: first, previous work found that the precise timing of phase asynchrony may influence how swaps errors arise ^26^ and second, splitting trials into reciprocal-swap and single-swap left an insufficient number of trials for PPI analysis.

The number of trials showing reciprocal-swap was in the range 7–81 across subjects (*M*=46.50, *SD*=18.51), while for single-swap was in the range 15–123 (*M*=69.88, *SD*=31.70), demonstrating that swaps are not necessarily the result of reciprocal/symmetric feature exchanges. For the alpha-FSV over trials, we observed a negative cluster (alpha-FSV higher in single-swap than reciprocal-swap trials), extending over latencies between stimulus presentation and the beginning of the delay period (∼0–0.740 s; Fig. 6a). This cluster included 63 sensors, mostly in frontal and sensorimotor sites, over both hemispheres (Cohen’s *d* in the range 0.071–0.617 across sensors; Fig. 6b). The cluster-based permutation test revealed a significant difference between swap types in alpha-FSV over trials (*p_perm_*=0.0060). We did not observe significant differences later in the delay for alpha-FSV higher in reciprocal-swap than single-swap trials. Further, we did not find any statistically significant difference between swap types in the alpha-FSV over time (lowest *p_perm_*=0.443 among clusters). When we repeated the analysis in source-space, we observed a trend towards increased alpha-FSV over trials for single-swap compared to reciprocal-swap trials, in areas of the lateral and medial prefrontal cortex, temporal pole–TP, and angular gyrus–AG, in particular in the contralateral hemisphere, which is in line with the sensor-space results; however, these effects did not reach statistical significance with a rigorous statistical threshold (positive cluster extending approximately over latencies 0–1.844 s; *p_perm_*=0.0919; see Fig. 6c–d).

**Fig. 6.**
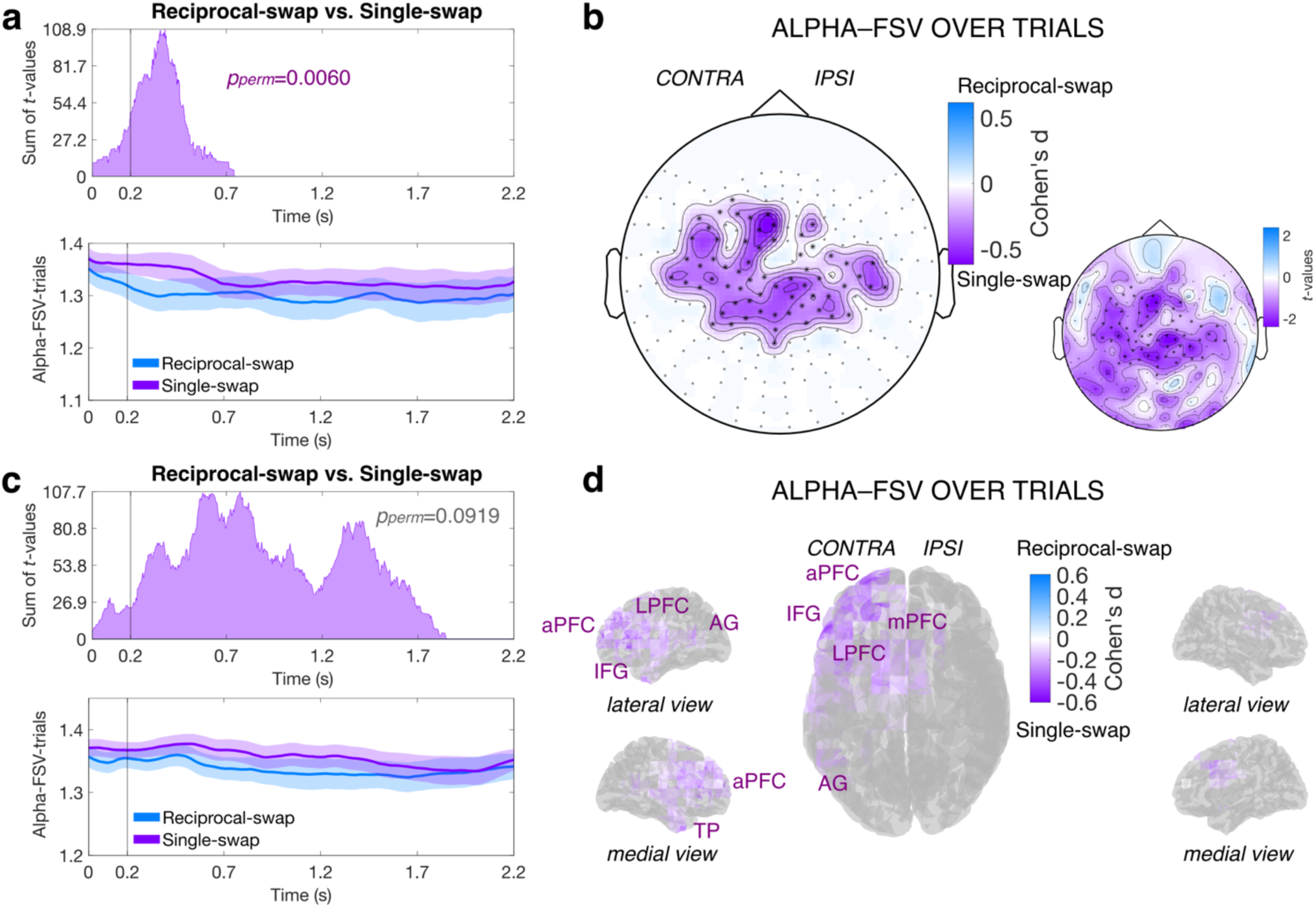
Differences in alpha-FSV over trials between reciprocal-swap and single-swap trials. **a** Time distribution of the sum of alpha-FSV differences across MEG sensors (*t*-values; top), and time course of the average alpha-FSV across sensors of the observed cluster for each condition (bottom). The shadings represent the standard error of the mean. **b** Topography plot with superimposed effect sizes of alpha-FSV differences between reciprocal-swap and single-swap trials, for each MEG sensor of the observed cluster (*t*-values are shown on the smaller topography plot on the right). **c** Time distribution of the sum of alpha-FSV differences across source points in source-space (*t*-values; top), and time course of the average alpha-FSV across source points of the cluster with the lowest *p*-value from the permutation test (*p_perm_*=0.0919) for each condition (bottom). The shadings represent the standard error of the mean. **d** Glass brain with superimposed effect sizes of differences between reciprocal-swap and single-swap trials.

### 2.5. Increased alpha power during WM maintenance

To confirm that the phase effects shown in this study are not due to simpler, lower-level differences in power spectral properties, we performed the following control analysis. We performed a time-varying spectral analysis and compared power estimates between trial types (see section 4.9). We found no statistically significant power differences between HP targets and swaps (the lowest *p*-value from the permutation test among the identified clusters was *p_perm_*=0.297), between HP targets and LP targets (lowest *p_perm_*=0.119), or between LP targets and swaps (lowest *p_perm_*=0.792). This suggests that the observed PPI and FSV results cannot be explained simply by lower-level differences in spectral properties across trial types. However, we found a task-related increase in alpha power (∼10 Hz) for all trial types, which was prominent during the memory delay (Fig. 7). To assess task-evoked changes in power at different frequencies, we estimated a measure of relative power change (RPC) with respect to baseline, by subtracting and normalizing the time-varying power estimates against the baseline average (pre-trial fixation period), separately for each trial type and for each MEG sensor (see section 4.9). The average RPC across all sensors showed an alpha power increase during memory retention across the different trial types (Fig. 7a). The observed alpha-band power increases were most pronounced in parietal-occipital MEG sensors, both contralateral and ipsilateral to stimulus presentation (Fig. 7b). Despite the lack of significant differences in power between trial types, these results suggest that alpha activity and synchrony tend to increase during the task, especially during WM maintenance.

**Fig. 7.**
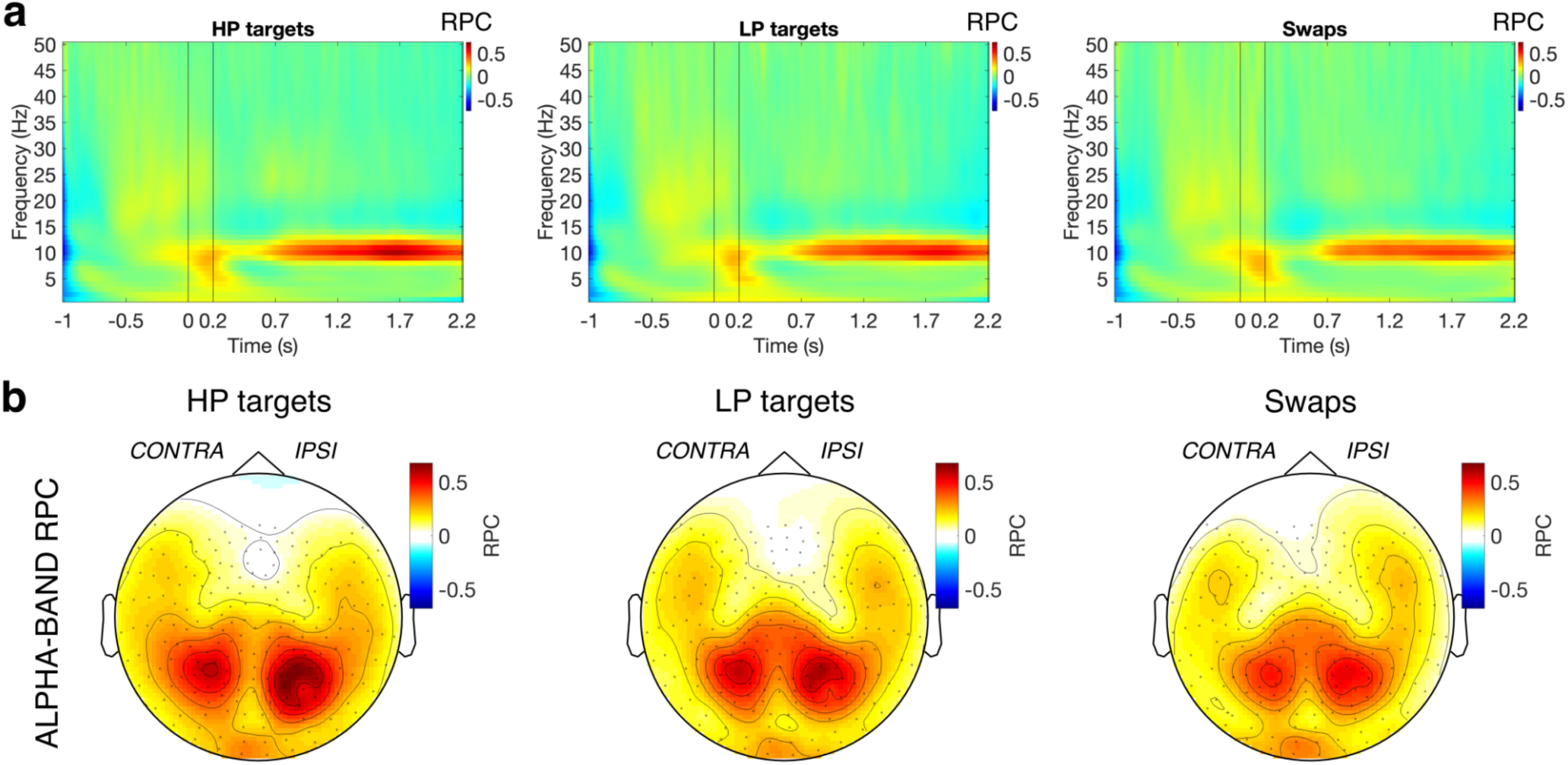
Relative power change (RPC) and alpha power increase during WM maintenance. **a** Time-frequency RPC in HP targets (left), LP targets (middle), and swaps (right). Each plot shows the grand-average RPC across all MEG sensors and subjects. Positive values are represented in red and indicate task-evoked increases in power (synchronization), while negative values in blue indicate task-evoked decreases (desynchronization). On the time axis: 0 s represents the onset of stimulus presentation and 0.2 s represents the onset of the memory delay. **b** Topography plot with superimposed the RPC in the alpha band (8–12 Hz), averaged across the delay period. The plot is shown for HP targets (left), LP targets (middle), and swaps (right). Positive values are represented in red and indicate task-evoked increases in alpha power during WM maintenance.

## 3. Discussion

Previous models have proposed alternative neural processes for how the binding of information between properties or features that belong to an object is accomplished in WM ^10,11^. An influential class of theories and models proposed that feature binding is accomplished via the phase synchronization of signals of neurons (phase coding) that store the different feature values corresponding to the memorized item ^13,14,18,24–26^. Here, we tested the central neurophysiological prediction of these models. We found that the correct feature binding is supported by stable phase-locked oscillatory activity at alpha frequencies, while swaps (binding errors in memory) are characterized by reduced phase preservation during WM maintenance, with sources localized in the hemisphere contralateral to stimulus presentation. This reduction did not generalize to other types of errors, suggesting that alpha phase inconsistencies are a hallmark of swaps. These results align with the idea that representational stability in WM exists along a continuum, rather than as a binary distinction between accurate and erroneous binding ^11,43^. Swaps may emerge from degradation or temporal incoherence in phase coding that falls below the threshold necessary for correct feature conjunction ^26^. An independent analysis found that these phase inconsistencies arose from increased phase coding variability. We localized the increase in phase coding variability for swaps in contralateral premotor, motor, parietal, and visual cortical areas. Thus, we provide convergent evidence that, in swaps, the sources of reduced neural phase synchrony and higher phase coding variability are in contralateral areas. Together our results suggest that feature binding in WM is accomplished through alpha phase coding dynamics, which supports both detailed computational models of feature integration in WM ^26^ and more general theories of time-based binding for cognitive flexibility ^44^. More broadly, this work represents a demonstration of how mechanistic predictions generated by biophysically-realistic models can be tested using human neuroimaging. Modeling studies suggest that phase coding is supported by the competition between different memories, and may result from lateral inhibition between neural populations selective for different features. In particular, Barbosa et al. proposed a model where the maintenance of conjunctions is accomplished through phase coding between two coupled one-dimensional attractor networks ^26^. In the model, oscillatory dynamics emerge within each network through the local interplay of fast recurrent excitation and slower feedback inhibition. In turn, different populations active in each network (representing different colors or different locations) oscillate out-of-phase based on the competition between the activity bumps, accomplished by gamma-aminobutyric acid (GABA)-mediated lateral inhibition between them. Finally, feature conjunctions are established via weak non-specific excitatory connections between the two attractor networks, whereby pairs of bumps oscillating in each network synchronize selectively. Several empirical observations are consistent with this mechanistic account. Strong feedback inhibition is a prominent feature of cortical networks ^45^, and it strongly controls their activity fluctuations ^46^. Also, brain activity is known to be dominated by an alternation in the firing of different neuronal populations ^47^, and the ‘gating by inhibition hypothesis’ has proposed that alpha activity in sensory regions implements a mechanism of pulsed inhibition that silences neuronal firing ^48,49^, mediating the observed alternation in neuronal firing. Our results suggest that such alternating, phase coding mechanism supports feature binding during WM maintenance: stable oscillatory target dynamics result in high-performance reports of cued items, while noisy fluctuations produce abrupt alpha phase instabilities, inducing shifts to non-target dynamics that ultimately result in swaps.

We provide converging evidence that sources of reduced phase synchrony in swaps and increased phase coding variability are localized in areas contralateral to the visual stimuli, with overlapping effects observed in both sensor-space topographies and source-space reconstructions. However, the time windows of these effects appear disjoint across the two analyses—Phase Preservation Index (PPI) and Frequency Sliding Variability (FSV). While it may be tempting to ascribe distinct neurobiological roles to this temporal dissociation—such as attributing early PPI effects to encoding and late FSV effects to maintenance or retrieval—we argue that these differences likely reflect methodological constraints rather than true functional segregation. Specifically, the discrepancy can be explained by the inherent properties of the two measures, Phase Preservation Index–PPI and frequency sliding variability–FSV over trials, as well as the intrinsic limitations of the cluster-based permutation approach. Cluster-based permutation testing does not allow to precisely determine the exact timing of the observed effects, but it only provides an approximation of the extent of effects ^35,36^. In terms of the measures, PPI estimates exhibit a rapid asymptotic decay over time from the reference point ^26,33^, which, in our case, is the onset of the memory delay—making them sensitive primarily to early differences. This was confirmed in a control analysis using a later reference point, which showed a similar spatial/frequency pattern of alpha-band PPI differences near the new reference. In contrast, FSV is not subject to this temporal bias. Alpha-FSV over trials does seem to differentiate between swaps and HP targets not only late in the delay but also early, especially in the sensor-space analysis (see Fig. 5a, section 2.3). However, this was not significantly detected by the cluster analysis. Taking into account these methodological limitations, our data is consistent with the view that quick, rapid phase changes occurring at any time during the delay are responsible for producing swap errors ^26^.

Both phase-locked alpha activity (indexed by PPI) and alpha-frequency sliding variability (FSV) revealed a consistent gradient across trial types: high-performance (HP) targets showed the most stable phase dynamics, swaps the least, and low-performance (LP) targets fell in between. Specifically, LP trials exhibited reduced phase-locked alpha activity and increased phase coding variability relative to HP targets, yet remained more stable and less variable than swaps. The observed gradual trend from HP to swaps, with LP in between, even in the absence of statistically significant differences between LP and swaps, suggests a graded relationship between phase stability and the fidelity of feature binding. Within an attractor dynamics framework, HP trials may reflect strong convergence on well-formed attractors, while LP trials hover near the edge of these basins—sufficient to maintain partial fidelity but vulnerable to drift or interference. Swaps, by contrast, may result from representations that fail to stabilize within the appropriate attractor basin altogether, or that fall into nearby but incorrect attractors due to increased phase noise or temporal instability. While not conclusive, this provides valuable exploratory insight and aligns with the notion of a continuum of representational stability ^11,43^, suggesting that swaps may emerge when representations fall below a stability threshold, not because of complete failure, but due to partial degradation or temporal incoherence in phase-based maintenance ^26^.

While our results highlight the importance of alpha activity, oscillatory activity in the theta (4–8 Hz) frequency band also plays a critical role in the neural mechanisms underlying WM maintenance, and the two are thought to reflect distinct yet complementary functional processes ^50^. Theta oscillations, especially over midline frontal areas, are consistently associated with the active maintenance and coordination of information in WM. Frontal-midline theta power increases with memory load and task difficulty, suggesting a role in executive control and the allocation of cognitive resources ^51,52^. Additionally, theta-band synchrony is thought to support long-range communication between prefrontal and posterior brain regions, facilitating the integration of distributed mnemonic representations ^53,54^. Alpha oscillations, particularly over parieto-occipital regions, have been linked to functional inhibition and the suppression of task-irrelevant sensory input. During WM maintenance, increased alpha power is thought to reduce interference from external stimuli by disengaging sensory processing pathways ^48,55^. This inhibitory function supports the maintenance of internal representations by protecting them from irrelevant or distracting input ^56^. Together, alpha and theta rhythms orchestrate a balance between inhibitory gating and top-down control, enabling robust and efficient maintenance of information over short durations. Their coordinated dynamics reflect a broader network-level organization crucial for supporting WM processes ^57,58^. Our results confirmed that alpha power increases during the maintenance phase of WM tasks, particularly in parietal and occipital cortices in both hemispheres. However, we localized the effects of phase-locked alpha activity specifically in regions contralateral to stimulus presentation, suggesting that alpha phase functions as a temporal organizing mechanism, enabling the integration of distinct feature dimensions into coherent memory representations. Our findings provide the first direct evidence that alpha phase coding supports feature binding in WM, confirming the predictions of the biologically-plausible, binding-by-synchrony attractor model ^26^.

Our findings that alpha phase coding supports feature binding in WM agree broadly with previous studies indicating that mid-frequency alpha and beta oscillations play a central role in the control over WM ^55,59–64^. Nonhuman primate WM studies using invasive recordings in the monkeys’ prefrontal cortex, for example, have found an inverse relationship between gamma (∼40–120 Hz) and beta power (∼20–35 Hz), mediated by oscillatory bursts ^65–67^. During WM maintenance these anti-correlated beta and gamma dynamics have been found to underlie a reduction in spiking variability ^68^, which supports the notion that mnemonic representations are maintained through mechanisms of phase coding ^18,23^. While conceptually consistent with our observations, these findings differ from our own both in the regions (prefrontal cortex vs. sensorimotor areas) and frequency band (beta vs. alpha). A potential reconciliation comes from a recent fMRI study that used longitudinal training to show that item-selective mnemonic representations become detectable in the prefrontal cortex over long-term learning ^69^. Hence, one possibility is that our observed phase coding mechanisms supporting the storage of mnemonic representations, may spread from posterior alpha to frontal beta over the course of training or, more in general, when task-specific categories, associations, and rules are learned.

The fact that phase synchrony disruptions were localized to sensory and motor cortices is consistent with the ‘sensorimotor recruitment’ model of WM, which states that visual information is maintained in the same stimulus-selective regions that are responsible for perceiving that information ^2,70–73^, as well as with the ‘distributed system view’ of WM ^74,75^, which states that WM storage is distributed across multiple brain regions, among which both visual and parietal cortices play an important role in storing visual information during WM maintenance ^76,77^. This is in line with the proposal that occipital and posterior parietal cortices are involved in the process of binding during visual WM, in particular in the maintenance of the bindings, which is supported by previous studies showing that delay period activity in the intraparietal sulcus–IPS predicts behavioral and neural correlates of binding at recall ^78,79^, as well as by the report that the influence of location-context binding on the representation of stimulus features is strongest in occipital cortex ^80^. Causal evidence for the involvement of these regions in information storage and in the binding process during WM has also been provided using noninvasive neurostimulation techniques ^81–83^. The model proposed by Barbosa and colleagues relies on two ring-attractor networks, one for each feature space (color and spatial location), thus it explicitly simulates the independent storage of individual features constituting the item ^26^, in line with the increasing evidence that different features are stored in independent brain systems (different cortical areas) ^11,84^. In our task, storage for color and spatial location may occur in distinct neural populations. For example, the colors of the items may be stored in color sensitive areas in visual cortex, while their spatial locations may be integrated into salience maps by premotor/motor and parietal areas, by storing them in systems that represent them not only as perceptual information but also as targets for future motor responses during the report period. Our data is consistent with the view that sudden phase instabilities in either system produce abrupt changes in the phase relationship between the two systems, changing the correct conjunctions between features and producing binding errors.

The presence of binding errors has been examined also in cued-recall tasks where two non-spatial features were used as the cue and report features ^85–87^; however, spatial location has been recognized as having a privileged role in feature binding and in the generation of swaps compared to non-spatial features ^29,87^. Previous studies have shown that the likelihood of swaps depends on the feature employed as the memory cue, and the frequency of swaps is higher when spatial location is the report feature compared to when it is the cue feature ^88–90^. In the present study, we employed color as the cue feature and spatial location as the report feature. While the number of swaps would be lower when using location as cue feature and color as report feature, we expect that our results would be similar in terms of modulations of alpha phase dynamics. Future work should explicitly evaluate this scenario to confirm this hypothesis. Also, the mechanistic conditions for this asymmetry should be addressed in computational models, as they currently only consider symmetric feature networks ^26^. We see at least two possibilities to break this symmetry: having stronger within-region noise for the color networks or asymmetric readouts.

The origin of swaps in WM remains a topic of ongoing debate, and the relationship of swaps to feature binding is still controversial. Some studies have proposed that swaps derive from an informed guessing strategy, rather than being true binding errors in memory ^91,92^. Other studies, however, have countered this account by providing evidence that swaps are better explained by cue-dimension variability ^90^ or by demonstrating that it is possible to reconstruct a mnemonic representation of the swapped location but not of the cued location, which suggests that swaps involve the active maintenance of a non-target memory item ^40^. One study investigated the cause of swaps by manipulating the variability with which either cue or report features are encoded, and provided supporting evidence for the hypothesis that swaps can be fully accounted for by variability in memory for the cue feature ^90^. By using behavioral modeling, this study showed that the ‘neural binding model’ ^29^—a model that incorporates feature binding within memory for single features—is superior to different variants of the ‘interference model’ ^93,94^. Interference models imply the presence of both cue-dependent and cue-independent swaps, which are failures in binding that are independent of individual features and are unrelated to cue-feature similarity. Swap errors in these models are predominantly attributed to cue-independent processes ^94^. However, cue-independent swaps did not occur in the study; moreover, the results were not consistent with a strategic response to forgotten items ^90^.

Recent studies have also shown that swaps are reflected in neural activity even before the response period, providing further evidence against a strategic guessing account of swaps. One neuroimaging study using fMRI demonstrated that swaps are preceded by the active maintenance of non-target memory items, rather than by the spontaneous production of random guesses at the response stage ^40^. Another study analyzed the neural population recordings from two rhesus macaques performing a color WM task ^95^. Using simultaneous recordings from posterior parietal cortex, lateral frontal cortex, motor cortex, posterior inferotemporal cortex, and visual area V4, the study provided evidence that swaps are induced during the selection of information. These ‘selection errors’ were associated with the neural activity in the second delay period (after retro cue), and they were also found in the second delay period of a separate ‘prospective’ task, in which the animal had to attend to one stimulus from a set of two visual stimuli. Because the neural data were modeled using all units with a non-zero firing rate, regardless of the cortical area from which they were recorded, it remains unclear whether neural processes in specific areas (e.g., executive control and motor regions) are the main source of these swaps emerging during the selection process from WM.

Furthermore, by examining the dynamics of mnemonic information on single trials obtained from large, local populations of lateral prefrontal neurons recorded in monkeys performing a WM task, one study found that cue-specific spiking activity does not persist during the memory delay, but is instead intermittent, alternating between coordinated ‘on’ and ‘off’ states ^96^. These states correspond to robust coding of the mnemonic representations and complete lapses in such coding, respectively. Although limited to the lateral prefrontal cortex, the study suggests that these transitions occur in coordination across large neuronal populations. In the previous study ^95^, mnemonic information about the non-target memory item may persist more in the spiking activity of neuronal populations during the delay associated with the selection process, when irrelevant information is discarded and relevant information is gated out from WM, becoming accessible for motor planning before the response period. Thus, the resulting swaps might be more detectable through classification from the fitted behavioral model and could be attributed to the selection process. However, they may be less consistently detected during the memory delay, in line with the notion of coordinated, intermittent coding of the mnemonic representations ^96^, as well as with the oscillatory dynamics during WM, which emerge from bump competition in the ring attractor models of spiking neural networks ^26^. As a matter of fact, the authors found that, although this was not consistent across the two monkeys, some swaps emerged while multiple stimuli were remembered during the first delay period of the WM task, prior to the retro cue ^95^. These findings show an association between swaps and misbound representations, suggesting that some swaps can indeed be directly attributed to neural misbinding in memory. Our work adds to this literature by demonstrating increased variability in alpha phase coding for swaps during WM maintenance in distributed sensorimotor areas. This supports the notion that swaps are not mere response guesses but true binding errors in memory that occur prior to the initiation of a response.

Beyond demonstrating that swaps are associated with disruptions in phase coding dynamics, we found that swaps can arise through both a reciprocal/symmetric and a unidirectional feature exchange. When we compared these two types of swaps, we found higher alpha phase coding variability in single-swap (single non-target report) than reciprocal-swap trials (at least two non-target reports). These effects were observed predominantly at latencies between the stimulus presentation and the beginning of the delay period, over medial, frontal, and sensorimotor sensors. These results suggest that single-swap trials may also capture situations in which a wrong association between visual features is encoded during the initial perception and encoding of the items’ features into WM, even before memory retention ^38,39^, and/or in the early moments of WM maintenance. Based on the topology of the differences, these swaps may derive from phase coding variability in a different network of brain regions, suggesting that they may thus derive from a different process. A caveat of this analysis is, however, that the number of trials was limited for the two swap types. Future studies are required to demonstrate whether distinct sources of swaps can be characterized on the basis of modulations in phase coding variability.

The current work investigates how phase coding dynamics contribute to maintaining bound representations during WM retention, with task design and analyses specifically optimized to examine neural activity during the delay period. However, it is important to acknowledge that swap errors may also originate from failures during earlier or later stages of processing—specifically, during encoding or recall. Prior studies have shown that binding errors can arise when attention is divided at the time of encoding, leading to incorrect associations between features such as color and location ^38,41^. Similarly, disruptions caused by eye movements or shifts in spatial attention have been found to produce feature-binding errors, suggesting that attentional stability is critical not only for maintenance but also for establishing accurate feature conjunctions ^39^. Moreover, work by Schneegans and Bays ^11^ emphasizes that feature binding in visual WM involves dynamic interactions across neural populations throughout all phases of memory processing. Recent evidence also indicates that neural signatures of swap errors are detectable at recall, pointing to the possibility of retrieval-based misbinding ^40^. Notably, the attractor model ^26^ also recognizes that swap errors can originate from multiple sources: in addition to ‘memory swaps’, it identifies ‘attentional swaps’ (i.e., when an incorrect association is encoded during the encoding period) and ‘decoding swaps’ (i.e., when the correct association is encoded and maintained during the memory period, but retrieval fails). These findings collectively suggest that swap errors likely result from a combination of failures in encoding, maintenance, and recall, highlighting the need for an integrated framework that captures binding dynamics across the full temporal course of WM. Future work will be essential to disentangle the distinct neural mechanisms underlying encoding-, maintenance-, and recall-related binding failures, in order to develop a comprehensive account of how and when feature-binding errors emerge in WM.

Overall, our results provide empirical support for time-based binding theories and highlight alpha-phase stability as a potential biomarker for binding integrity. Future work might build on these findings by applying single-trial classification methods or causal manipulations (e.g., frequency-specific transcranial magnetic stimulation) to further establish the functional role of alpha synchronization in feature binding.

## 4. Methods

### 4.1. Subjects

This study was performed in compliance with the Declaration of Helsinki on “Medical Research Involving Human Subjects”, and was approved by the IRB ethics committee at the New York University (NYU) Abu Dhabi. Healthy human subjects from the community of NYU Abu Dhabi were recruited to participate in the study. Subjects participated as paid volunteers (50 AED/h) and gave written informed consent prior to the experimental sessions. All subjects had normal or corrected-to-normal vision. In a preliminary behavioral session, subjects performed the WM task presented in Fig. 1. Only subjects who made a ‘reciprocal-swap’ (see section 4.2) in at least 5% of the trials were invited for a following MEG session. Twenty-six subjects participated in the MEG session (age range 19–37 years, *M*=24.33, *SD*=4.57; 4 female; all right-handed). Previous papers reporting Phase Preservation Index have ranged from 6 to 12 subjects and typically do not report effect size ^97–100^; we based our sample size on an estimated effect size of 0.5 and power of 0.8. Both behavioral and MEG sessions were characterized by the same experimental design and procedures (see below).

### 4.2. Stimuli, design, and procedures

Visual stimuli consisted of 3 different items (colored circles), each of which measured 0.55 degrees of visual angle (DVA) in diameter, presented at an eccentricity of 4.5 DVA (Fig. 1). Stimulus items were presented on a gray background for 0.2 s, followed by a retention interval (delay period) during which the screen was blank (2 s). A white fixation point (0.2 DVA) was presented at the center of the screen during both stimulus presentation and delay period. In a pre-fixation period (1 s before the beginning of each trial), the fixation point changed color to red when the subjects broke fixation. Trials in which the subjects broke fixation were removed from successive analyses.

Subjects were asked to remember the 3 stimulus items, and to hold in mind their colors and locations during the delay period. After memory delay, subjects were asked to report the location of each of the items (report period), which were sequentially cued by presenting their color at the center of the screen. Subjects had no control over the order in which the items were cued during the report period. During the report, a white circle (same size and eccentricity of the items) was also presented on the screen. In each trial, the spatial position of this report circle was randomly drawn from a discrete uniform distribution of integer values between 1° and 360° of polar angle. Subjects were asked to respond by adjusting the position of the report circle by using a rotating dial, and pressing a button on a response box to confirm their response. The report period was self-paced and was followed by a feedback period (1 s), showing the cued items ringed by white circles and the subjects’ responses ringed by black circles. After the feedback, a self-paced inter-trial interval was used to allow the subjects to control their blinking. Subjects had to press a button to move onto the next trial.

Each subject performed 500 trials of the WM task in one session, divided into 10 blocks (50 trials each). During stimulus presentation, the items were always presented on one single hemifield (5 blocks on the left, 5 blocks on the right; randomly ordered), never across hemifields. In each trial, the spatial location of each stimulus item was randomly sampled from a space that depended on the hemifield used for that block, obtained by excluding locations that were 10° of polar angle away from the vertical meridian, and using a minimum gap of 15° of polar angle. This was done to prevent physical overlap between items and to avoid swaps purely induced by misperception of spatial location. The color of each item was selected randomly from a color wheel of 180 color segments, using a minimum gap of 15 color segments between items to avoid swaps that were purely induced by colors misperception. These color segments were drawn from a circle in CIELAB color space, with radius of 59°, and centered at *L\**=54, *a\**=18, *b\**=−8, where the *L\** axis represents lightness, and the *a\** and *b\** axes represent chromaticity (red-green and yellow-blue components, respectively) ^28,101^. This approach is commonly used to create a psychophysically uniform color space ^102,103^.

Typically, in the presence of swap errors, responses would be clustered around non-target features and the histogram of response deviations should show a central peak, around deviations of zero ^31^. However, the use of a minimum distance between the feature values in each trial (like in our experiment) may obscure the presence of such a central peak. To correct for these effects, we computed the response deviations using a permutation of the non-target feature values, specified relative to the target feature, across trials ^29^. This method removes the signatures of actual swaps, while still capturing the effects of the minimum feature distance, which provides an expected histogram of non-target deviations in the absence of swaps. We then computed the difference between the histogram of actual non-target deviations and the histogram of expected non-target deviations, obtaining a distribution of non-target response deviations corrected for the effects of minimum feature distance. This distribution displays a central peak if swaps are present, while it would be approximately uniform if swaps are absent.

We used a maximum likelihood approach to assign to each response a likelihood of being a binding error (non-target) or a target response. The approach was based on a mixture model that comprised the probability of correctly reporting the target item, the probability of incorrectly reporting a non-target item, and the probability of responding randomly (guess) ^30^ (https://www.paulbays.com/toolbox/). This procedure allowed categorizing each trial on the basis of the subjects’ responses: trials with a probability of correctly reporting the target item greater than 0.95 in all three responses were considered high-performance (HP) targets, while those with a probability of incorrectly reporting a non-target item greater than 0.70 in at least one response were considered swaps. The remaining trials were considered low-performance (LP) reports. After MEG data preprocessing (see section 4.3), the number of trials across subjects was in the range 16–439 for HP targets (*M*=92.81, *SD*=88.06), 22–196 for swaps (*M*=116.38, *SD*=47.79), and 44–328 for LP trials (*M*=198.73, *SD*=64.03). For a specific analysis on swaps (see section 2.4), these were further divided into reciprocal-swaps (at least two non-target reports) and single-swaps (single non-target report). The number of trials for the two swap types was in the range 7–81 (*M*=46.50, *SD*=18.51) and 15–123 (*M*=69.88, *SD*=31.70) across subjects, respectively.

For each subject, we refitted the data with the mixture model separately for the trials of each type (HP trials, LP trials, and swaps) and derived the concentration parameter κ of the von Mises distribution, which describes the response variability for that trial type. The concentration parameter provides a measure that is the reciprocal of the dispersion of response errors (i.e., the inverse of the circular standard deviation); thus, it provides a measure of the precision of WM. We compared the concentration parameter between trial types using a paired samples *t*-test (*df*=25).

### 4.3. MEG data acquisition and preprocessing

The locations of the marker coils on each subject, as well as fiducial locations (including nasion–NAS, left and right Pre-Auricular points–LPA/RPA, left and right PreFrontal–LPF/RPF), were recorded before MEG data acquisition. The shape of the subject’s head was also recorded using a Polhemus dual source handheld FastSCAN-II system (Colchester, VT, United States of America), collecting between 2,500 and 4,500 points per subject. MEG data were acquired continuously using a 208-channel axial gradiometer system (Kanazawa Institute of Technology, Kanazawa, Japan), with sampling frequency of 1,000 Hz, and applying an online low-pass filter (cutoff frequency 200 Hz). During recording, the subject was lying in the supine position, while performing the WM task. The visual stimuli were generated using MATLAB (The MathWorks Inc., Natick, USA) and projected onto a screen positioned at 85 cm from the face of the subject.

We applied a noise reduction procedure to the continuous MEG data using the rotated spectral Principal Component Analysis (rsPCA), as implemented in the MEG laboratory software MEG160 (Yokogawa Electric Corporation, Tokyo, Japan), with block width of 5000 and 30 shifts, by using eight magnetometer reference channels that were located away from the subject’s head. We used the FastICA algorithm implemented in MNE-Python ^104^ to decompose the MEG data by independent component analysis (ICA), and we removed the components identified as eye blinks or heartbeat artifacts from the data. We defined trials using the time epoch from 1 s before stimulus onset (pre-fixation period) to 2.7 s after stimulus onset. We then applied temporal demean and low-pass filtering (cutoff frequency 140 Hz) procedures to the data using FieldTrip ^105^ (http://www.ru.nl/neuroimaging/fieldtrip/). Noisy trials and channels were automatically identified (high variance) and rejected by visual inspection. We rejected trials showing residual ocular or muscular artifacts by visual inspection. We interpolated the rejected channels using the average of all their neighbors.

As a final step, we switched the MEG channels between the right and left sides in all trials where the stimulus was presented in the left hemifield. This pooling procedure was performed to combine trials from both left-hemifield and right-hemifield stimulus presentations, treating them as if the stimulus presentation had always appeared in the right hemifield. The contralateral/ipsilateral pooling was justified by our findings, which showed no significant differences in absolute mean error between trials with left-hemifield and right-hemifield stimulus presentations, nor in the proportion of left/right trials across different trial types. These were tested using paired samples *t*-tests (*df*=25). We found no significant difference in absolute mean error (*t*(25)=1.1910, *p*=0.2393) between trials with left-hemifield (*M*=25.3°, *SD*=5.6°, range 13.4°–37.2°) and right-hemifield (*M*=23.5°, *SD*=5.1°, range 13.9°–33.3°) stimulus presentations. Further, we found no significant difference in the proportion of left/right trials across the different trial types: HP trials (*t*(25)=-0.7055, *p*=0.4871; left: *M*=45.50, *SD*=44.45, range 6–220; right: *M*=47.31, *SD*=44.57, range 10–219), LP trials (*t*(25)=-0.7554, *p*=0.4571; left: *M*=98.54, *SD*=33.61, range 15–156; right: *M*=100.19, *SD*=31.36, range 29–172), and swaps (*t*(25)=1.3411, *p*=0.1920; left: *M*=60.00, *SD*=26.41, range 12–110; right: *M*=56.38, *SD*=23.22, range 10–95). We note that the pooling analysis improved sensitivity to changes in phase contralateral to stimulus presentation while decreasing sensitivity to activity that is lateralized regardless of stimulus presentation. As our interest was in exploring changes in phase relationships tied to working memory storage-related activity, which displays consistent contralateral organization ^106–108^, pooling maximized our ability to identify signals of interest related to our main hypothesis.

### 4.4. Statistics

We performed all statistical comparisons with a cluster-based permutation approach ^35^, using a two-tailed dependent *t*-test (*df*=25, *p*<0.05, alpha level distributed over both tails by multiplying the *p*-values with a factor of 2 prior to thresholding), 1,000 permutations, and *p_perm_*<0.05 for the permutation test. We used the same settings in every comparison, unless specified otherwise, while the space for clustering in each analysis is specified in the sections below.

In each comparison between trial types, we estimated the effect sizes of the differences using Cohen’s *d* ^109^, after averaging the estimates separately per trial type, either over the time points and frequencies identified by the significant cluster (Phase Preservation Index–PPI analysis; see section 4.5), or simply over time points (frequency sliding variability–FSV analyses; see section 4.6–4.7).

### 4.5. Phase Preservation Index

We used the Phase Preservation Index–PPI ^33^ to estimate the consistency over trials of time-frequency phase differences with respect to a reference phase, separately for each trial type (HP, LP, and swaps). Specifically, PPI measures the intertrial consistency in phase differences as a function of time (*t*), with respect to a reference time point (*t_ref_*), at a specific frequency of interest *f_0_*:

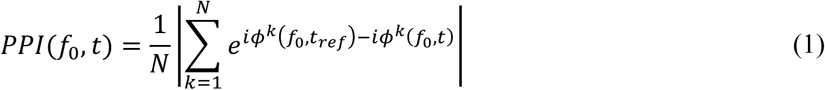

For each trial (*k*=*1,…, N*) we derived the instantaneous phase *Φ^k^(f_0_,t)* using a Morlet wavelet transform (central frequency parameter *ω_0_*=6; zero-padding to solve edge effects problem) ^110–112^, in each MEG sensor. The frequency of interest *f_0_* varied in the range 1–50 Hz (1 Hz steps). We used the delay onset as the reference time point (*t_ref_*=0.2 s) for PPI estimation, following the previous work on the attractor model ^26^. In a separate analysis, we repeated the PPI estimation using the middle of the delay as the reference time (*t_ref_*=1.2 s), to provide a more complete picture of the timing of the observed PPI effects.

Since PPI is sensitive to the number of trials, this was balanced between trial types (HP targets and swaps) at the level of each subject. We performed this procedure by: (i) considering 100 combinations of trials subsets in the condition with more trials, to match the number in the condition with fewer trials; (ii) estimating PPI for each subset; and (iii) computing the median of PPI estimates across subsets. We estimated PPI only once in the condition with fewer trials. We then compared PPI between HP targets and swaps using a cluster-based permutation approach (see section 4.4), over all time points in the delay period (0.2–2.2 s), frequencies (1–50 Hz), and MEG sensors. Here, we defined neighboring sensors based on their 3D geometrical distance (max distance of 25 mm for defining neighbors).

To confirm the presence of within-trial phase synchronization, we derived ‘PPI corrected’ estimates as the difference between the PPI estimates obtained from the previous analysis and those obtained from a randomization procedure. Here, we randomly shuffled the signals over time, for each MEG sensor and trial independently. We then repeated the PPI estimation on time-shuffled data (temporally uncorrelated) ^33^, including the procedure to equate the number of trials between conditions, as described above. Finally, for each subject we computed the difference between PPI estimates from the original data and those obtained from time-shuffled data.

### 4.6. Frequency sliding (instantaneous frequency)

We employed the frequency sliding–FS ^37^ to capture the single-trial temporal fluctuations in oscillation peak frequency. FS estimation relies on five core steps: (i) extract the phase angle time series at a certain frequency (Fig. 8a); (ii) unwrap the phase angle time series (Fig. 8a); (iii) compute the first order derivative over time (measure in rad); (iv) scale the measure by multiplying by the sampling frequency and dividing by 2π (measure converted into Hz; Fig. 8b); (v) apply a median filter to attenuate neurophysiological noise spikes (Fig. 8b). We employed a Morlet wavelet transform to extract the phase angle time series (step i), using *ω_0_*=6 and zero-padding ^110–112^. We used an order of 10 and a maximum window size of 0.1 s for the median filter settings (step v), which reassigned each time point to be the median of a distribution made from surrounding points ^37^.

**Fig. 8.**
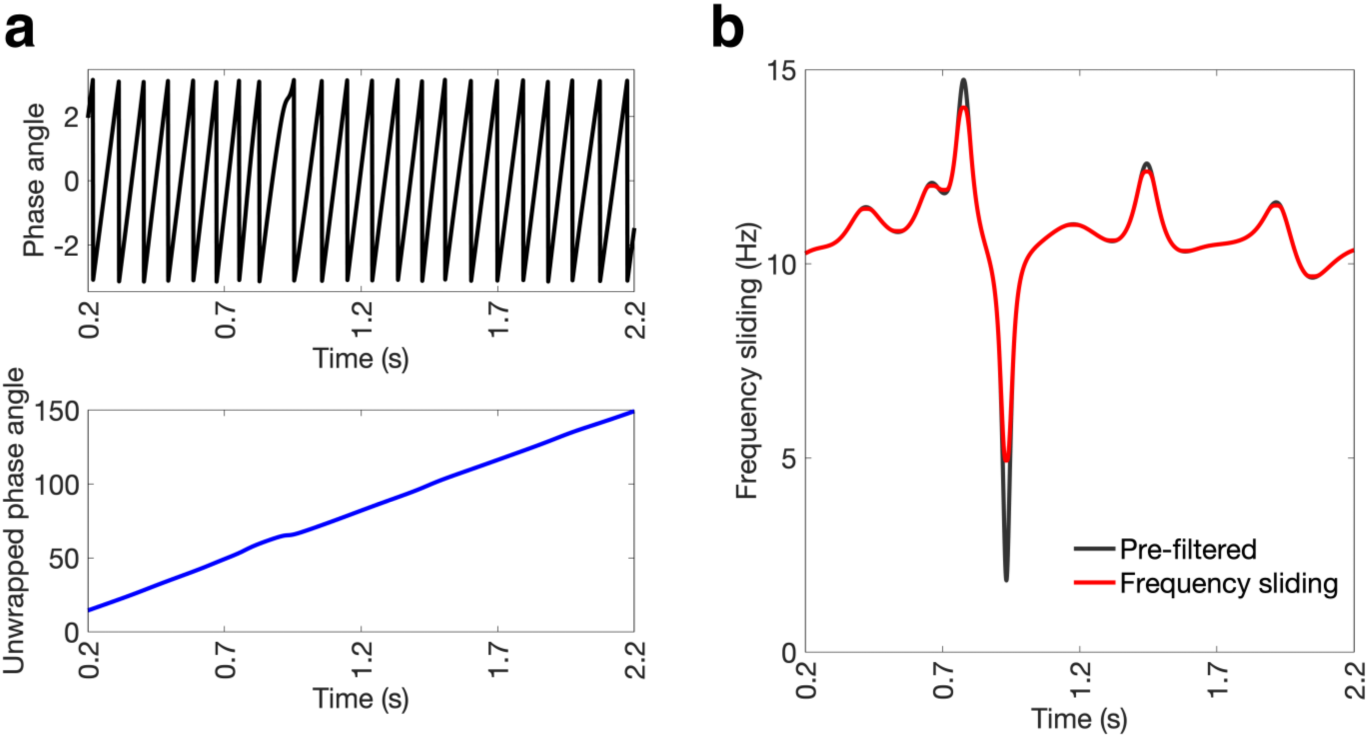
Estimation of frequency sliding–FS (instantaneous frequency). **a** Example of phase angle time series at 10 Hz, before (top, in black) and after (bottom, in blue) the unwrapping procedure. **b** Frequency sliding pre-filtered (black) and after median filtering (red) for the phase angle time series shown in panel a.

To derive measures of FS variability (FSV) over time and over trials, we first derived single-trial FS estimates at 10 Hz, separately for each trial type; then, we estimated the median between trials of the standard deviation of FS over time points (alpha-FSV over time) and the median between time points of the standard deviation of FS over trials (alpha-FSV over trials) on a sliding window of 0.4 s (i.e., four cycles of the 10 Hz oscillation), with maximum overlap in time. We finally compared the two measures of alpha-FSV between trial types using a cluster-based permutation approach (see section 4.4), over time points during the stimulus presentation and in the delay period (0–2.2 s) and sensors, where neighboring sensors were defined based on their 3D geometrical distance (max distance of 25 mm for defining neighbors).

In a separate control analysis, we compared alpha-FSV over trials and alpha-FSV over time in swaps. For each subject, we first computed the average of each alpha-FSV measure over time points in the delay period and sensors from the significant cluster, obtained from the previous analyses (20 contralateral, parietal-occipital sensors; see Fig. 5b). Then we compared the average alpha-FSV over trials and alpha-FSV over time using a paired samples *t*-test (*df*=25).

### 4.7. Source reconstruction

The solution of the MEG forward problem requires models of the head volume conduction, the sensors, and the cortical sources. To derive these models, we estimated a transformation matrix (rotation and translation) through an automatic procedure that aligned the five fiducials (NAS, LPA, RPA, LPF, and RPF) between subject-specific Polhemus head shape and markers. We then used the transformation matrix to align the sensors’ locations to the head shape. We used the boundary element method (BEM) template provided in FieldTrip ^105^ for the head volume conduction. We coregistered the template head model with the head shape, and successively refined the transformation using a method that relies on fitting spheres to the head shapes (which allows deriving translation and global scaling). We computed the head volume conduction based on the refined template head models, by creating a single-shell model on the basis of the brain compartment, and taking its inner shell. As a final step, we manually realigned the head model and head shape, and the registration was also checked manually. The source model was constructed using template brain coordinates (12.5 mm resolution; 10 mm outward shift; 1,499 equivalent current dipoles), and then transformed into head coordinates. We computed the subject-specific lead field matrix considering a forward operator with unconstrained orientation, where each source point was modeled as three orthogonal equivalent current dipoles.

Once the lead field matrix was computed, we solved the MEG inverse problem using the linearly constrained minimum variance (LCMV) beamformer ^113^. We obtained an estimate of sensor-space covariance matrix from the 1 s pre-fixation period. We extracted source-reconstructed time series from all source points using a method based on singular value decomposition ^114^, which allows to derive a single-trial time series for each source point that best explains the variability in dipole orientations and signal strengths across trials for that point. After source reconstruction, we derived measures of PPI, single-trial FS and FSV at 10 Hz for each source point, following the same procedures used in the sensor-space analyses (see sections 4.5–4.6). We then compared PPI and the two measures of FSV between HP targets and swaps using a cluster-based permutation approach (see section 4.4), over time points either in the delay period only (0.2–2.2 s; for PPI) or in both stimulus presentation and delay period (0–2.2 s; for alpha-FSV), and source points in source-space. We defined neighboring points based on their distance in MNI space (max distance of 15 mm for defining neighbors).

### 4.8. Computational model

We used the attractor model of spiking neural networks described by Barbosa et al. ^26^ to generate simulated field potentials from two networks implementing phase-encoded feature binding in working memory, in order to test the ability of FS to detect synchronization changes corresponding to swap trials. In brief, we built two identical networks of excitatory and inhibitory integrate-and-fire spiking neurons, such that within each network neurons were disposed on a ring and the connection strength of NMDA and AMPA receptors (for excitatory connections) or GABA_A_ receptors (for inhibitory connections) between any two neurons was defined by a Gaussian function of their arc distance on the ring (standard deviation, or footprint of the connectivity, σ). Strong recurrent excitation and larger footprint for inhibitory than excitatory connections facilitate tuned persistent activity (a bump) after brief stimulation of a subset of neurons. In turn, when two bumps are sustained in one ring network, broader inhibitory footprints impose out-of-phase synchronization of the two bumps (Fig. 4c–d). Connections between excitatory neurons in one ring and neurons in the other ring were instead non-specific, allowing for the arbitrary synchronization of bump pairs across the two networks. More specific details of the network implementation can be found in Barbosa et al. ^26^, or in the code repository (https://github.com/comptelab/binding). We computed the extracellularly recorded field potential for each of the network by applying a Gaussian filter (σ=5 ms) centered on each spike from neurons in one half of the network (simulating biases in representing the visual space), to obtain a time series for each network in each trial (Fig. 4c–d).

### 4.9. Power analysis

We performed a spectral analysis separately for each trial type (HP, LP, and swaps) using the Morlet wavelet transform with a central frequency parameter of *ω_0_*=6 ^111,112^, to derive time-varying power estimates at frequencies from 1–50 Hz, in 1 Hz increments. The estimation was performed over a time interval that included the pre-fixation period (1 s before the beginning of each trial), stimulus presentation (0.2 s), and the delay period (2 s). We applied zero-padding to mitigate edge effects ^110^. Power spectra were compared between trial types using a cluster-based permutation approach (see section 4.4), over all time points in the interval (-1.0 to 2.2 s), frequencies (1–50 Hz), and MEG sensors. Neighboring sensors were defined based on their 3D geometrical distance, with a maximum distance of 25 mm for defining neighbors.

To investigate whether there were any systematic task-evoked increases (synchronization) or decreases (desynchronization) in power at specific frequencies, we computed a measure of relative power change (RPC) with respect to baseline ^115^. RPC was estimated separately for each trial type as the time-varying change in power relative to baseline:

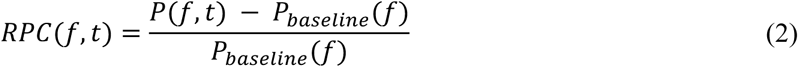

Where *P(f,t)* was the power at time *t* and frequency *f*, and *P_baseline_(f)* was obtained by averaging the power spectra over time frames in a baseline period (-1.0–0 s prestimulus, i.e., the pre-fixation interval), in each MEG.

### 4.10. Data availability

Analysis scripts, behavioral data, and preprocessed MEG data are available on the Open Science Framework (https://osf.io/s5mcy/; https://osf.io/37hb6/). The computational model can be obtained from https://github.com/comptelab/binding. Supplemental data can be requested via email to the lead author.

## Acknowledgements

This work was supported by the Swiss National Science Foundation (Grants P2FRP3_195083 and P500PB_214404 to M.F.P.), the National Institutes of Health (Grant R01MH063901 to M.D.), and the NYU Research Enhancement Fund and the NYUAD Center for Brain and Health, funded by Tamkeen under NYUAD Research Institute (grant CG012 to K.K.S.). A.C. and J.B. acknowledge support from the Spanish Ministry of Science, Competitiveness and Universities co-funded by the European Regional Development Fund (Refs: BFU 2015-65318-R, RTI2018-094190-B-I00). J.B. was supported by the Bial Foundation (ref: 356/19). This research was conducted using resources provided by the High Performance Computing Cluster and the Core Technology Platform at NYUAD. We would like to thank Vahan Babushkin for assistance with piloting and technical support.

## Author Contributions

M.F.P. and K.K.S. conceptualized the study. K.K.S. and A.T. developed the experimental paradigm. A.T. collected the data. M.F.P. and A.S. preprocessed the data. M.F.P., A.S., A.T., J.B., A.C., and K.K.S. developed the analytical framework. M.F.P. and K.K.S. wrote the manuscript. A.S., J.B., A.C., and M.D. provided feedback and edited the manuscript.

## Supplementary Information

**Fig. S1.**
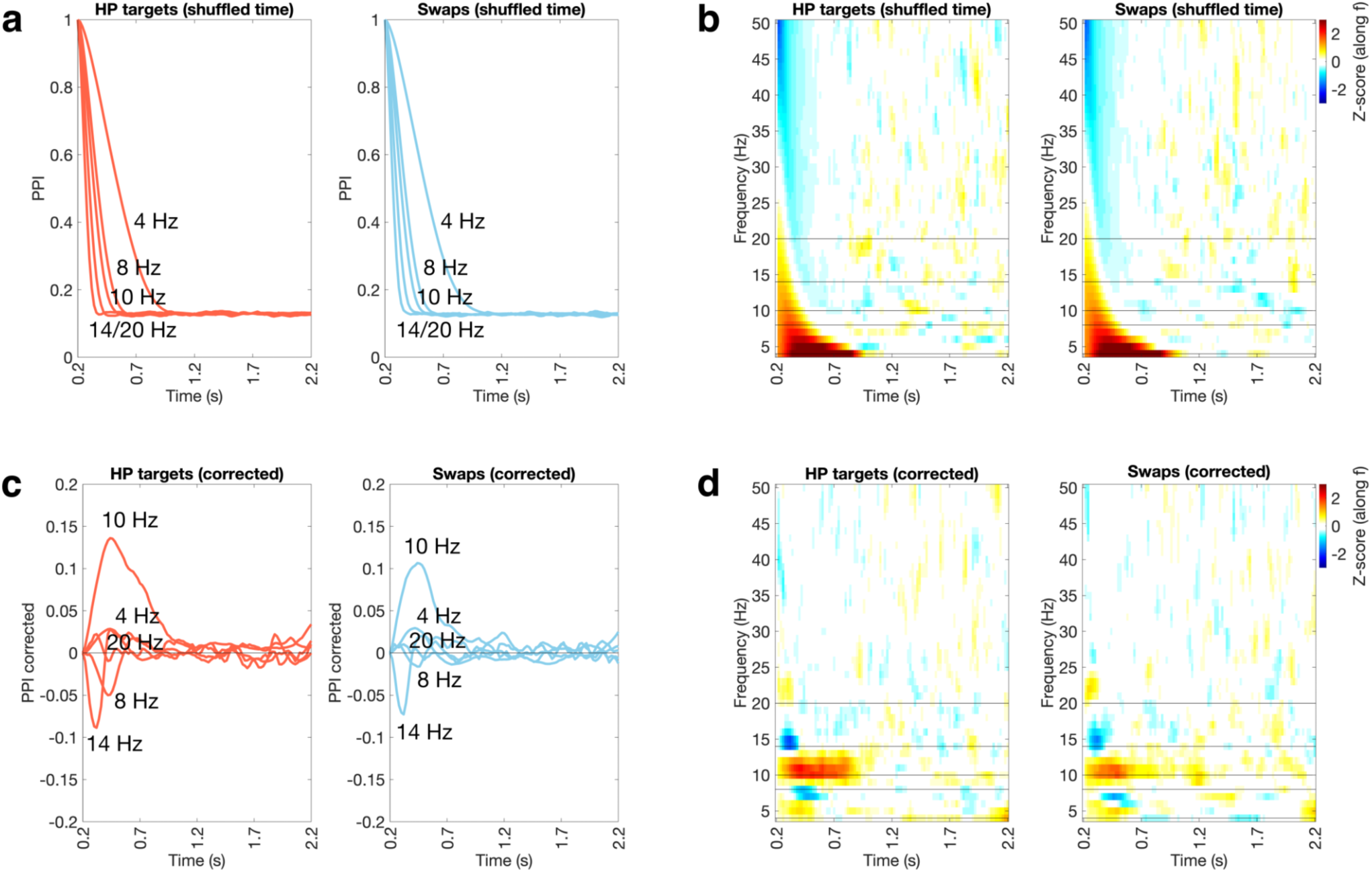
PPI estimates between HP targets and swaps. **a** Time course of PPI obtained from shuffling the signals over time in HP targets (left) and swaps (right) at different frequencies, highlighted by the black horizontal lines in the next panel. **b** PPI (shuffled signals over time): z-scores along frequencies for each time point in the delay, in HP targets (left) and swaps (right). **c** Time course of PPI corrected in HP targets (left) and swaps (right) at different frequencies, highlighted by the black horizontal lines in the next panel. **d** PPI corrected: z-scores along frequencies for each time point in the delay, in HP targets (left) and swaps (right).

**Fig. S2.**
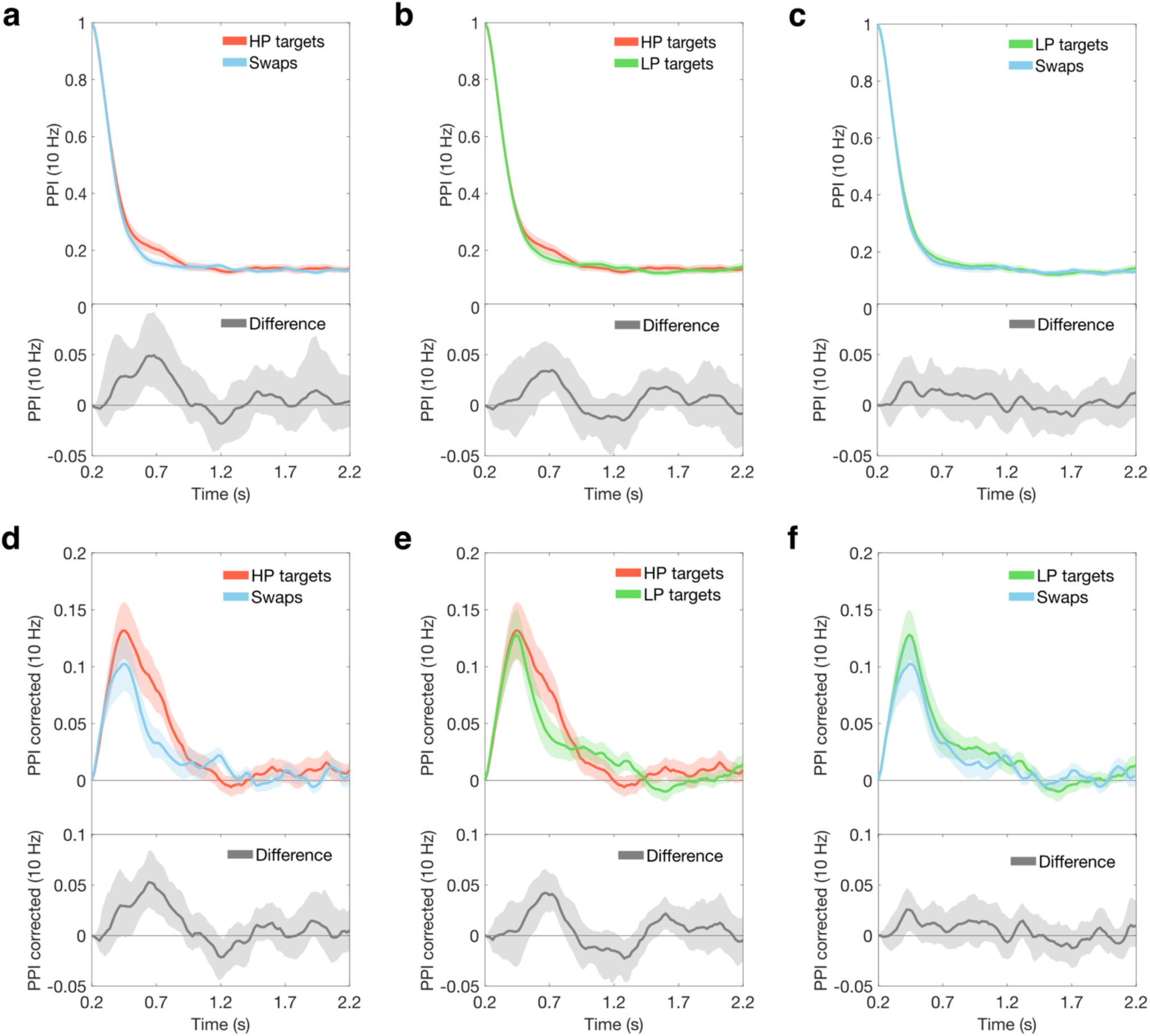
PPI estimates across MEG sensors of the cluster in the observed data, for the different trial types. **a** Time course of PPI at 10 Hz for HP targets (red) and swaps (blue), together with the difference between the two (gray). **b** Time course of PPI at 10 Hz for HP targets (red) and LP targets (green), together with the difference between the two (gray). **c** Time course of PPI at 10 Hz for LP targets (green) and swaps (blue), together with the difference between the two (gray). **d** Time course of PPI corrected at 10 Hz for HP targets (red) and swaps (blue), together with the difference between the two (gray). **e** Time course of PPI corrected at 10 Hz for HP targets (red) and LP targets (green), together with the difference between the two (gray). **f** Time course of PPI corrected at 10 Hz for LP targets (green) and swaps (blue), together with the difference between the two (gray). In each panel, the colored shadings (top) represent the standard error of the mean, while the gray shading (difference; bottom) represents 95% confidence intervals (CIs). CIs were estimated using the bias-corrected and accelerated method on a bootstrap distribution of across subjects differences, obtained by resampling with replacement 10,000 times.

**Fig. S3.**
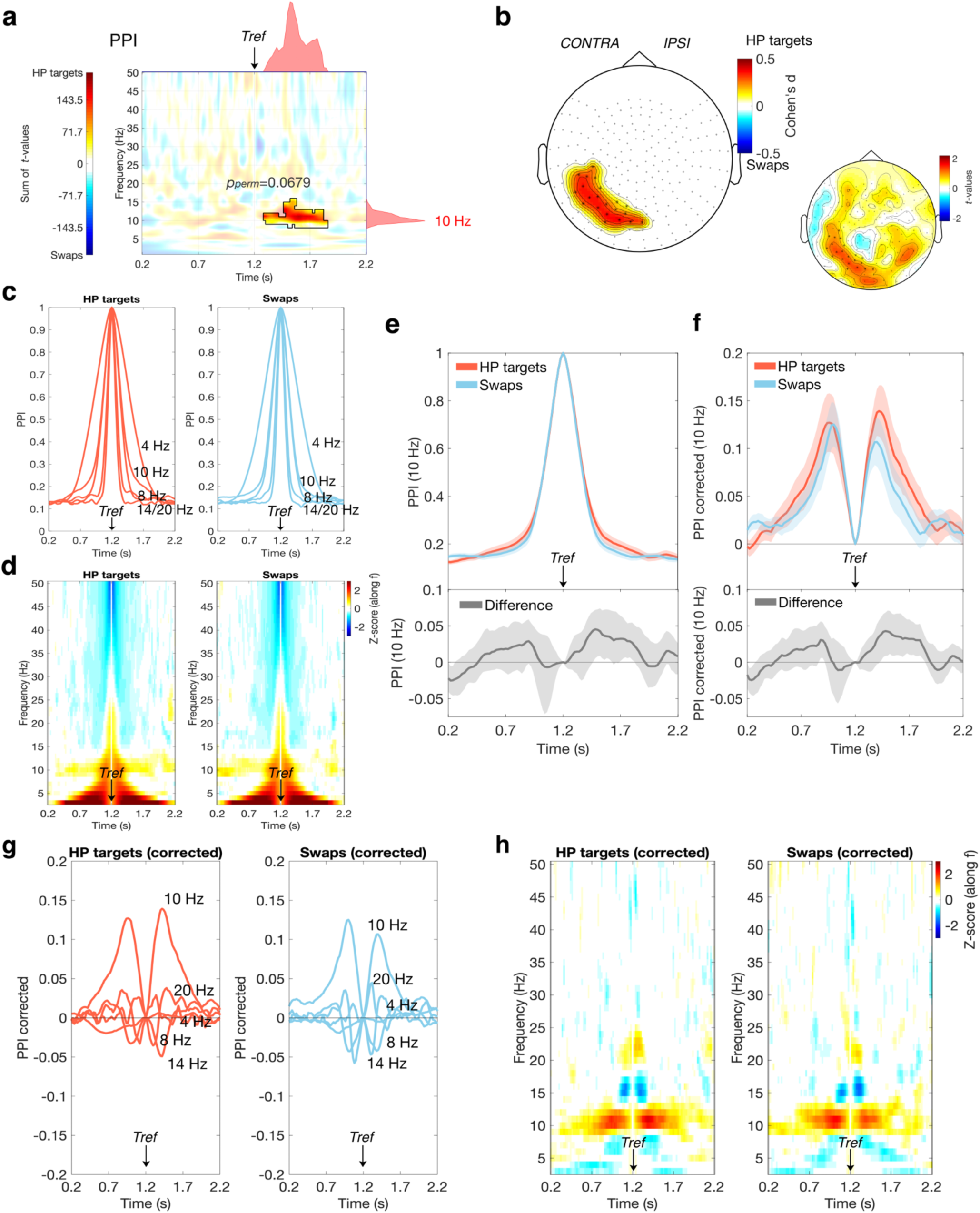
PPI differences between HP targets and swaps: analysis using the middle of the delay as the reference time point (*t_ref_*=1.2 s) for PPI estimation. a Time-frequency distribution of the sum of PPI differences across MEG sensors (*t*-values). The black contour highlights the positive cluster found in the observed data. The marginal plots on the right and on top represent respectively the time-collapsed frequency distribution and frequency-collapsed time distribution of the differences between trial types. **b** Topography plot with superimposed effect sizes of PPI differences between HP targets and swaps, for each MEG sensor of the observed cluster (*t*-values are shown on the smaller topography plot on the right). The positive cluster did not reach statistical significance in the cluster-based permutations testing (*p_perm_*=0.0679) **c** Time course of PPI in HP targets (left) and swaps (right) at different frequencies (as in Fig. 3). **d** Z-scores along frequencies for each time point in the delay, in HP targets (left) and swaps (right). **e** Time course of PPI estimates at 10 Hz for HP targets (red) and swaps (blue), together with the difference between the two (gray). **f** Time course of PPI corrected at 10 Hz for HP targets (red) and swaps (blue), together with the difference between the two (gray). In e–f, the red and blue shadings (top) represent the standard error of the mean, while the gray shading (difference; bottom) represents 95% confidence intervals (CIs). CIs were estimated using the bias-corrected and accelerated method on a bootstrap distribution of across subjects differences, obtained by resampling with replacement 10,000 times. **g–h** Same as in c–d, but for PPI corrected.

**Fig. S4.**
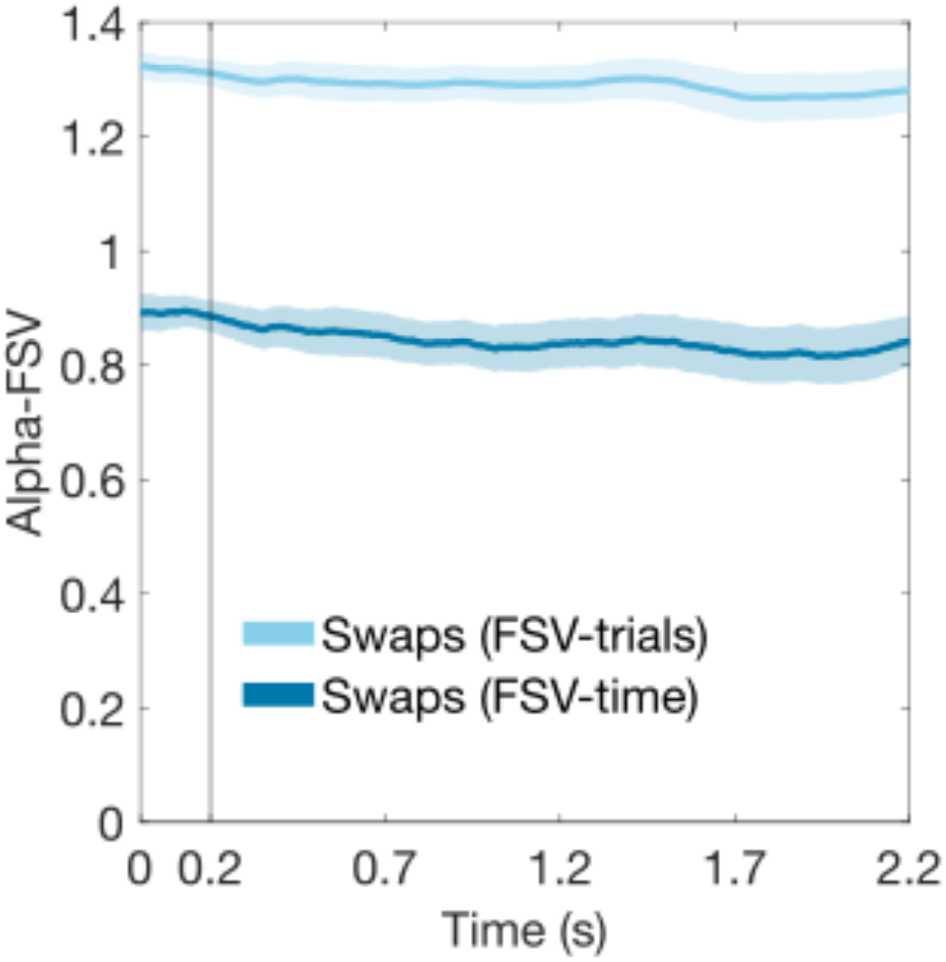
Measures of alpha-FSV in swaps. The time course of the average alpha-FSV across sensors of the cluster found in the data (see Fig. 5b) is shown for alpha-FSV over trials (light blue) and alpha-FSV over time (darker blue). The shadings represent the standard error of the mean.

**Fig. S5.**
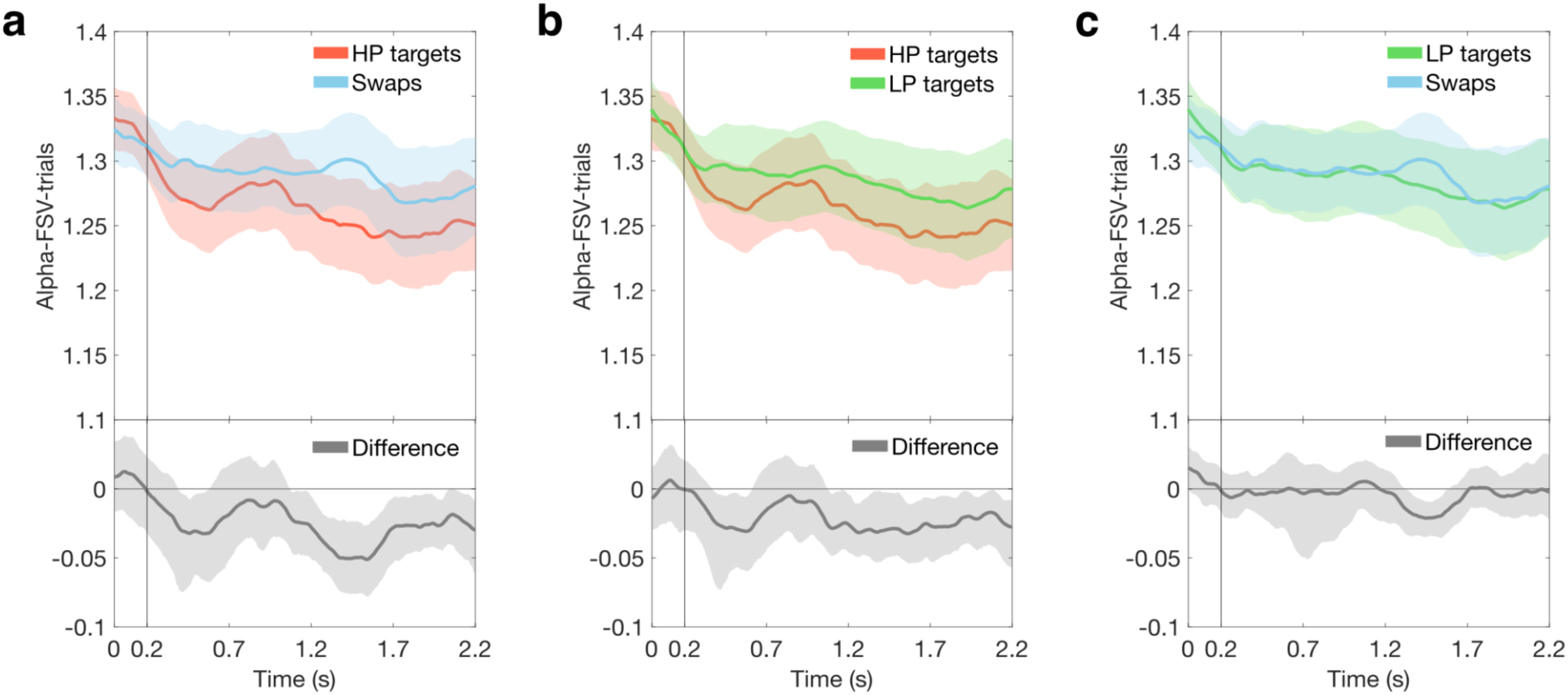
Alpha-FSV over trials across MEG sensors of the cluster in the observed data, for the different trial types. **a** Time course of alpha-FSV over trials for HP targets (red) and swaps (blue), together with the difference between the two (gray). **b** Time course of alpha-FSV over trials for HP targets (red) and LP targets (green), together with the difference between the two (gray). **c** Time course of alpha-FSV over trials for LP targets (green) and swaps (blue), together with the difference between the two (gray). In each panel, the colored shadings (top) represent the standard error of the mean, while the gray shading (difference; bottom) represents 95% confidence intervals (CIs). CIs were estimated using the bias-corrected and accelerated method on a bootstrap distribution of across subjects differences, obtained by resampling with replacement 10,000 times.

